# *Chromosome-Level Genome of Simulium vittatum* Links Black Fly Cytogenetics to Genome Organization and Evolution

**DOI:** 10.64898/2026.09.08.750124

**Authors:** Erika Nishiduka, Tom Hill, Stephen Lu, Brian Bonilla, Elverson Melo, Lilian Caesar, Gabriel da Luz Wallau, Paola Valenzuela-Leon, Lucas Tirloni, Osvaldo Marinotti, Eric Calvo

## Abstract

Black flies (Diptera: Simuliidae) are important vectors of pathogens affecting human and animal health, yet the absence of a chromosome-level nuclear reference genome has constrained molecular and evolutionary studies of the family. Here, we present the first chromosome-level nuclear genome and a developmental transcriptomic resource for the long-established IS-7 laboratory lineage of *Simulium vittatum*. Combining Oxford Nanopore long-read sequencing with Hi-C scaffolding, we assembled a 340.4-Mb genome, with 99.1% of the assembly resolved into three chromosome-length scaffolds (N50=104.8 Mb, BUSCO completeness 93.9%), consistent with the known 2n = 6 karyotype. Using the historically mapped molecular landmarks SVAT and SVEP, we assigned the two arms of chromosome III as IIIS and IIIL, respectively, linking sequence coordinates to the classical polytene chromosome map. Repetitive DNA comprises 46.48% of the assembly, including 29.68% unclassified repeats, and annotation identified 14,732 protein-coding genes and 16,417 transcripts. This reference genome connects classical black fly cytogenetics with sequence-level analyses of genome organization, structural variation, and gene content, addressing a major genomic gap within Culicomorpha and providing a foundation for comparative studies of chromosome evolution, hematophagy, and vector biology across Simuliidae.

## Introduction

Black flies (Diptera: Simuliidae) are hematophagous insects whose immature stages develop in flowing freshwater habitats (1). Females of many species are important pests and vectors of pathogens affecting humans, livestock, and wildlife (1,2). Black flies are the only known vectors of *Onchocerca volvulus*, the causative agent of human onchocerciasis. (river blindness), and transmit pathogens of veterinary importance, including *Leucocytozoon* parasites and vesicular stomatitis virus (VSV) (3–7). Beyond their role in pathogen transmission, black fly swarms can cause substantial nuisance and economic impacts through the blood-feeding activity of females on humans and livestock, with heavy biting associated with blood loss, irritation, stress, and reduced livestock productivity (1,2).

Black flies have an unusually extensive cytogenetic literature. Polytene chromosome polymorphisms, particularly paracentric inversions, have long been used to resolve cryptic taxa, characterize population structure, investigate sex linkage, and infer evolutionary relationships within Simuliidae (8–10). In black flies, sex determination is frequently associated with differentiated regions of otherwise homomorphic chromosomes rather than with highly differentiated heteromorphic sex chromosomes (10). This cytogenetic work has developed alongside physiological and molecular studies of hematophagy, including the characterization of salivary anticoagulants, vasomodulators, extracellular matrix-degrading enzymes, and other factors that facilitate blood feeding and host interaction (11–18).

The long-established IS-7 laboratory lineage of *Simulium vittatum*, a widespread North American species, is a particularly valuable experimental system because multiple lines of research have converged on this lineage. Founded from eggs collected near Cambridge, New York, in 1981, the colony has been maintained through successive generations under laboratory conditions, enabling studies of colony biology and reproduction (19,20), vector competence (20), salivary physiology and hematophagy (11–18), and chromosome polymorphism and sex linkage (20). Cytogenetic analyses have documented both sex-linked and autosomal inversions in IS-7 (20), and two genes encoding antihemostatic salivary factors, SVAT and SVEP, have been mapped directly to its polytene chromosomes (12). Together, its long-term maintenance, experimental tractability, detailed cytogenetic maps, and molecularly mapped loci provide a well-established framework for integrating a chromosome-level reference genome with existing cytogenetic knowledge.

Although draft nuclear genome assemblies of *Simulium* have recently begun to emerge (Hedtke *et. al*.; Research Square, https://doi.org/10.21203/rs.3.rs-10364888/v1, preprint), no chromosome-level nuclear reference genome has yet been reported for Simuliidae. Chromosome-level genomes are available for other Culicomorpha, including the ceratopogonid *Culicoides sonorensis* (2n = 6, 189 Mb) (21), the chironomids *Chironomus tentans* (2n = 8, 185 Mb) (22) and *Chironomus riparius* (2n = 8, 192 Mb) (23), and the mosquitoes *Anopheles gambiae* (2n = 6, 278 Mb) (24) and *Aedes aegypti* (2n = 6, 1.3 Gb) (25). The extensively characterized brachyceran *Drosophila melanogaster* (2n = 8, 143.7 Mb) (26) provides an additional, but substantially more distant, dipteran genomic reference. These lineages diverged from Simuliidae approximately 145 million years ago for ceratopogonid and chironomid lineages, approximately 185 million years ago for mosquitoes, and more than 220 million years ago for the Brachycera lineage represented by *D. melanogaster* (27–29). These evolutionary distances limit the extent to which existing reference genomes can resolve lineage-specific patterns of chromosome evolution in black flies. This limitation is relevant because chromosomal rearrangements are central to Simuliidae diversification, whereas mitochondrial haplotypes do not necessarily correspond to cytogenetically defined sibling groups (30).

Here, we present the first chromosome-level nuclear genome of *S. vittatum*, generated with Oxford Nanopore Technologies (ONT) long-read sequencing and Hi-C scaffolding and complemented by life-stage and post-blood-feeding transcriptomes. We used these resources to characterize chromosome organization, structural variation, repetitive DNA, candidate horizontally transferred genes, mitochondrial genome organization, and functional gene repertoires. The resulting reference genome integrates sequence coordinates with classical cytogenetic maps, providing a foundation for comparative genomic studies within Simuliidae and across Diptera.

## Methods

### Black fly colony and rearing

The colonized *S. vittatum* cytospecies IS-7 was obtained from the University of Georgia Black Fly Research and Source Center (Athens, GA, USA). The colony originated from eggs collected near Cambridge, New York, in 1981 and has been maintained continuously under laboratory conditions. Black flies were reared as previously described (19,20). Adult flies were maintained under standard colony conditions. For blood feeding, females were offered bovine whole blood in acid-citrate-dextrose (ACD, Lampire Biological Laboratories, Pipersville, PA, USA) supplemented with 100 mM ATP through artificial membrane feeders (NDS Technologies, Inc., Vineland, NJ, USA) covered with Parafilm® M (Amcor, Ann Arbor, MI, USA).

### Genomic DNA extraction, ONT library preparation, and sequencing

High-molecular-weight (HMW) genomic DNA was extracted from the whole body of a single adult male *Simulium vittatum* using the MagAttract HMW DNA Kit (Qiagen, cat. n°. 67563). DNA quality and fragment size distribution were assessed using a TapeStation 4200 system (Agilent, G2991BA), which indicated that most fragments exceeded 60 kb. DNA concentration was measured using the Qubit dsDNA BR assay kit (Invitrogen, Q32853). Approximately 287 ng of HMW DNA was sheared to a target size of 60 kb using a Megaruptor 3 instrument (Diagenode, B06010003) with the DNAFluid+ kit for viscous samples (Diagenode, E07020001) at speed 45 for one cycle. The sample was then concentrated to 50 μL with a Vacufuge Plus (Eppendorf, E-VPVCCS), and DNA quantity and size distribution were reassessed by Qubit and TapeStation, respectively.

Sequencing libraries were prepared from 228 ng of genomic DNA in a final volume of 48 μL using the Oxford Nanopore Ligation Sequencing Kit V14 (SQK-LSK114), following manufacturer’s protocol with two modifications. Long Fragment Buffer was used during adapter post-ligation cleanup to enrich for longer DNA fragments, and the final library elution was extended to 20 min at 37°C. The library was loaded onto an R10.4.1 flow cell (FLO-PRO114M) and sequenced on a PromethION 24 platform (Oxford Nanopore Technologies) (31) at the Frederick National Laboratory for Cancer Research (National Cancer Institute, Frederick, MD, USA).

### ONT read processing and quality control

Raw signal data were generated and stored in POD5 format using MinKNOW v24.05.14 and basecalled offline using Dorado v0.7.2 in high-accuracy mode. Reads with Q-scores > 9 were retained and exported in FASTQ format, then merged and decompressed for downstream analyses. Sequencing quality was summarized with pycoQC (31) and NanoStat (32), and potential contamination was evaluated using Kraken (33). The retained ONT long reads provided approximately 256x genome coverage.

### Hi-C library preparation and sequencing

Phase Genomics (Seattle, WA, USA) performed Hi-C sample preparation and library construction with the Proximo kit v4.5. Approximately 500 mg of whole male black flies were finely chopped before DNA crosslinking and proximity ligation. Subsequent steps followed the manufacturer’s protocol for preparation of paired-end libraries with Proximo reagents. Libraries were sequenced on an Illumina NovaSeq X Plus 25B flow cell.

### Genome assembly, Hi-C scaffolding, and assembly assessment

ONT long reads were assembled using HifiASM, in ONT ultralong integration mode, with default parameters including deduplication (34). Haplotype one was used for downstream analysis, while haplotype two was retained for haplotype comparisons. Hi-C data were used to scaffold and curate the assembly. Contact maps were visualized in Juicebox (35) and manually inspected for misjoins, inversions, and orientation errors; identified inconsistencies were corrected before the final chromosome-scale assembly was established. Assembly quality and completeness were evaluated with BUSCO version. 5.0.0 using the Diptera OrthoDB data v10 database (n=3,285 single-copy orthologs) and with QUAST (36).

### Comparative genome alignment and chromosome homology

Chromosome-level scaffolds were assigned to chromosomes I–III by comparing their relative sizes and centromere-to-telomere proportions with the established cytogenetic karyotype of *S. vittatum*. Putative centromere positions, inferred from chromosome-scale Hi-C interaction patterns and associated genomic features, were used to assess correspondence with the metacentric or submetacentric configurations described cytogenetically (12,20). For chromosome III, arm identity was further established using previously mapped salivary loci as molecular anchors: the SVAT and SVEP loci, cytogenetically localized to IIIS and IIIL, respectively, were mapped to the assembled chromosome and used to orient the short and long arms relative to the putative centromere (12). Together, chromosome size, centromere position, chromosome morphology, and locus-specific molecular anchors were used to relate the chromosome-level assembly to the classical polytene chromosome map. Whole genome alignments were generated with D-GENIES (37) to compare the *S. vittatum* assembly with the NCBI assemblies of *Aedes aegypti* (GCF_002204515.2), *Culicoides sonorensis* (GCA_047716325.1), *Culicoides brevitarsis* (GCF_036172545.1), and *Culex quinquefasciatus* (GCA_015732765.1).

### SGP-2 sequence validation

The annotated SGP-2 CDS was reconstructed directly from the chromosome II reference sequence using the CDS coordinates in the final GFF annotation. The two coding segments were extracted from the final genome assembly and reverse-complemented according to the minus-strand orientation. Exact junction sequences representing the genome-derived and reconstructed (+G) models were searched independently against the paired-end larval RNA-seq FASTQ files. Exact sequence matches were counted separately in R1 and R2. Representative junction-containing RNA reads were aligned to the final genome using BLASTN to confirm their genomic origin. The SGP-2 genomic region was additionally aligned against the original ONT-derived contig assembly to determine whether the sequence discrepancy arose during chromosome scaffolding.

### RNA sample collection and total RNA isolation

Samples were collected across the *S. vittatum* life cycle: late-stage larvae (third and fourth instars), mixed male and female pupae, non-blood-fed adult males and females, and blood-fed females at 0, 3, 8, 24, 48, and 72 h post-feeding (Table 1). Total RNA was extracted from pools of 3 to 5 individuals using TRIzol (Invitrogen, cat. n° 15596018) according to the manufacturer’s instructions. Each pooled sample was used to generate a single Illumina RNA-sequencing library. Because only one biological sample was available for each stage or condition, the RNA-seq data were used for genome annotation and descriptive expression profiling, and no formal statistical differential expression analyses were performed.

**Table 1:** Samples used for Simulium vittatum transcriptome sequencing.

| Sample condition | Group<br>Label | # Specimens<br>per sample | Collection time point |
| --- | --- | --- | --- |
| Larvae, third- and fourth-instar | L | 5 | 7 days after hatching |
| Pupae, late stage | P | 5 | 24-48 hours after pupation |
| Adult male, sugar-fed | M | 3 | 4-6 days after emergence |
| Adult female, sugar-fed | F | 3 | 4-6 days after emergence |
| Adult female, right after blood-feeding | F0 | 3 | 4-6 days after emergence |
| Adult female, 3 h post blood-feeding | F3 | 3 | 4-6 days after emergence |
| Adult female, 8 h post blood-feeding | F8 | 3 | 4-6 days after emergence |
| Adult female, 24 h post blood-feeding | F24 | 3 | 5-7 days after emergence |
| Adult female, 48 h post blood-feeding | F48 | 3 | 6-8 days after emergence |
| Adult female, 72 h post blood-feeding | F72 | 3 | 7-9 days after emergence |

### mRNA library preparation, quality control, and sequencing

Novogene (Sacramento, CA, USA) performed mRNA library preparation and sequencing. Total RNA quality was assessed by RNA Tapestation (Agilent Technologies Inc., California, USA) and quantified by AccuBlue® Broad Range RNA Quantitation assay (Biotium, California, USA). Samples were purified with mRNA Capture Module (Cat. # RK20340, ABclonal, Woburn, MA, USA), and paired-end (PE) libraries were generated according to the mRNA-seq Lib Prep kit Module for Illumina (RK20350, ABclonal, Woburn, MA, USA). Following fragmentation, first strand cDNA was synthesized using random hexamer primers, followed by second strand synthesis, end repair, A-tailing, adapter ligation, size selection, amplification, and purification. Final libraries were quantified with Qubit 2.0 (ThermoFisher, Massachusetts, USA), assessed with TapeStation HSD1000 ScreenTape (Agilent Technologies Inc., California, USA), and sequenced on an Illumina NovaSeq X Plus 25B platform (Illumina, California, USA).

### Transposable elements (TEs) discovery, classification, and manual curation

#### De novo identification and evidence-based classification of TEs

Repetitive sequences in the de novo assembly were identified with RepeatModeler (38), using -LTRStruct. RepeatClassifier assignments were retained for later curation. Satellite, Simple_repeat, rRNA, snRNA, tRNA, and Unknown (or not classified) consensuses <150 bp were removed. The remaining consensuses were clustered with cd-hit-est at ≥80% nucleotide identity and ≥80% coverage of the shorter sequence (-c 0.8 -aS 0.8 -G 0 -g 1 -b 500 -n 5).

The filtered library was characterized with PASTEC v2.0 (39), in the REPET framework, using BLASTN, BLASTX, and TBLASTX searches against Repbase 30.01 (40) Classification combined TE similarity, HMM protein profiles, species-specific exons, eukaryotic rDNA, tRNAs, and structural features (terminal repeats, poly(A) tails, tandem repeats, and open reading frames). PASTEC assignments were manually reviewed with RepeatClassifier annotations and structural, coding, and homology evidence. Final classifications were made case by case from TE-associated protein domains, similarity to characterized elements, structural features, and matches to host genes or other non-TE sequences, following general principles of manual TE curation (41).

Consensus sequences unresolved after PASTEC were further examined by homology and protein-domain searches. They were compared with the curated Dfam 3.9 HMM library using dfamscan.pl (E-value ≤1.10−5), and matches to Dfam families (42) were manually evaluated with existing evidence. For candidates with ORFs lacking informative Pfam annotations, we searched Swiss-Prot with Blastx and Conserved Domain Database (CDD) (43). with rpstblastn (BLAST+ v2.17.0; E-value ≤1.10−2). CDD assignments were processed with rpsbproc at 1.10−5, retaining specific and superfamily-level hits. TE-associated domains supported a TE origin, whereas cellular or other non-TE domains argued against it. Final classification combined these results with evidence from previous curation steps.

#### Manual refinement and extension of TE consensus sequences

Following the initial classification and curation steps, we manually refined and, when necessary, extended TE-supported consensus sequences following the Basic Protocol of Storer et al.(42). We first examined each consensus with TE-Aid against the genome assembly to assess genomic matches, internal structure, and completeness. We then recovered genomic copies with RepeatMasker (44) sing RMBlast in sensitive mode and organized them into family-specific alignments with generateSeedAlignments script.

Following *Storer et al*. (42), we iteratively refined the consensuses with alignAndCallConsensus script, using Kimura divergence to guide scoring-matrix selection and manually inspecting each multiple-sequence alignment. When aligned copies consistently supported sequence beyond the current boundaries, we incorporated flanking genomic sequence and extended the model. After each round, we reassessed the updated consensus with TE-Aid, including flanking regions, to evaluate boundaries and structural completeness. Refinement continued until the consensus stabilized or alignment and structural evidence indicated that the element boundaries had been reached.

#### Final TE library consolidation and genome-wide characterization

After consensus refinement, unresolved candidates were compared with curated TEs using BLASTN and TBLASTX. A custom script merged nonredundant BLASTN HSPs for each query-subject pair to estimate nucleotide identity and coverage without double-counting overlaps. Candidates satisfying the 80-80-80 criterion (at least 80% identity over at least 80% of their length, with an aligned region of at least 80 bp) were assigned to the same TE family. Redundant or fragmented consensuses, including shorter fragments of complete models, were collapsed, retaining the most complete curated sequence as the family representative.

BLASTN similarities below the family-level threshold supported more distant relationships and superfamily-level classification, while TBLASTX provided complementary evidence for relationships detectable at the translated-sequence level. These results were integrated with structural evidence from previous curation to resolve incomplete or unknown TE classifications. The final curated TE library was used to annotate the genome with RepeatMasker, using RMBlast in sensitive mode. RepeatMasker utilities were used to estimate total TE content and the relative representation of each superfamily. Divergence between genomic copies and their consensus sequences was calculated from RepeatMasker alignments and used as an approximate measure of relative TE age. Lower-divergence copies were considered relatively younger than more divergent copies. TE abundance across divergence classes was summarized to generate repeat landscapes for the major TE groups.

### Structural and protein-coding gene annotation

Repetitive regions from the *de novo* genome were annotated using RepeatMasker (44) (-gff - gcalc -s) and subsequently soft-masked using BEDTools (45) prior to gene annotation. RNA-seq reads were concatenated and quality filtered with Cutadapt (46) using the parameters --nextseq-trim=2 --trim-n -n 5 -O 5 -q 10,10 -m 35:3. Filtered reads were aligned to the soft-masked genome with STAR (47), and a genome-guided transcriptome assembly was generated with Trinity (48). The assembled transcriptome, soft-masked genome, *Aedes aegypti* proteome, the published *S. vittatum* sialome (11), and the UniProt protein database (49) were supplied as evidence to a first MAKER2 annotation run (50). Gene models from that run were used to train SNAP (51), and the species-specific prediction model was incorporated into a second MAKER2 iteration. In parallel, *ab initio* predictions were generated with Helixer through Galaxy (52–54). Single-exon models lacking transcript or protein support and models with annotation edit distance >0.3 were removed. MAKER2 and Helixer annotations were integrated with GFFcompare, and final coding sequences and predicted proteins were extracted with GFFread (55).

### Functional annotation

Predicted proteins were functionally annotated using an in-house rule-based program that parsed BLASTP and RPS-BLAST matches against the databases listed in Table 2. The program mapped terms from a curated vocabulary of approximately 450 keywords to broad functional classes and retained sequence identity and alignment coverage as supporting evidence.

**Table 2:** Protein databases used in the functional annotation of Simulium vittatum genome.

| Database Source |
| --- |
| CDD Database |
| MEROPS Database |
| NCBI Diptera Protein Database |
| PFAM Database |
| RefSeq Invertebrate Protein Database |
| RefSeq Protozoa Protein Database |
| RefSeq Mitochondrion Protein Database |
| Uniprot Protein Database |

Predicted proteins were assigned to broad functional classes after normalization of the original annotation categories. Functional identity and predicted localization were treated independently. Thus, transcripts carrying a secretion-associated annotation prefix were retained within their assigned biological class (e.g. protease, protease inhibitor, immunity, or metabolism), while secretion status was recorded separately from the annotation and SignalP results. Dedicated secreted-protein families without a more informative broad functional assignment (e.g. lipocalins, mucins, hormone-related proteins, and lipid-binding proteins) were retained in the Other.secreted.proteins class. Records containing protein-description text in place of a functional category were manually curated using available protein and conserved-domain annotations and plotted in R using ggplot2 package.

### Gene and transcript expression profiling

RNA-seek pipeline v1.9.0 (56) was used to quantify gene and transcript-level abundance, and to assess sample-level relationships by multidimensional scaling (MDS) and sample-to-sample expression heatmaps. Quality-control steps included FastQC, Preseq, Picard Tools, FastQ Screen (57), Kraken2 v2.0.8 (58), QualiMap (59), and RSeQC v2.6.4 (60). Adapters and low-quality sequences were removed with Cutadapt (46), and reads were aligned to the *S. vittatum* assembly with STAR v2.7.6a in per-sample two-pass basic mode (47). RSEM v1.3.3 (61) was used to calculate raw counts, transcripts per million (TPM), and fragments per kilobase per million reads (FPKM). Because each stage or condition was represented by a single pooled biological sample, expression analyses were descriptive and were not subjected to statistical differential-expression testing.

### Candidate sex-associated region analyses

Candidate sex-associated regions were evaluated by integrating several descriptive genomic analyses. Adult male and female transcript profiles were mapped to chromosomal coordinates using DEseq2 (62) to identify spatial clustering of genes with different transcript abundance; these comparisons were not tested for statistical differential expression. Reads assigned by Hifiasm to haplotype 1 or haplotype 2 were examined for haplotype-exclusive regions and summarized in 1-Mb windows. Long-read SNP variation and phasing were assessed with Clair3 and WhatsHap (63,64). BLASTP was used to identify *Drosophila melanogaster*, *Anopheles gambiae*, and *Aedes aegypti* orthologs and to examine correspondence with classical Muller elements. Because Muller elements are derived chromosomal units, these assignments were used as comparative homology evidence rather than as direct indicators of sex chromosomes. Paracentric inversions between haplotypes were evaluated with D-GENIES and DELLY in long-read mode (65). The chromosome-I haplotypes were also inspected specifically for evidence of the historically described IS-7 sex-linked inversion (20).

### Horizontal gene transfer analysis

Candidate horizontally transferred genes were identified by searching the predicted *S. vittatum* proteome against the NCBI non-redundant protein database (downloaded on November 10^th^, 2025) with DIAMOND v2.1.7.16 using --max-target-seqs 500, --min-score 50, and --outfmt 6. A custom script assigned NCBI Taxonomy annotations to BLAST hits (available at https://github.com/liliancaesarbio/alien_hgt_index). Hits were categorized as RECIPIENT (Insecta), GROUP (non-insect Metazoa), or OUTGROUP (non-Metazoa), and the E-value and bit score of the highest-scoring hit in each category were recorded for each protein. The Alien Index was calculated as AI = log(eG + 10⁻² □□) - log(eO + 10⁻² □□) (66), and the HGT index as hU = bO - bG (67). Proteins with AI > 30 and hU > 30 were retained as HGT candidates. To reduce false positives arising from contamination or symbiont-derived sequences, only candidates located on assembled genomic scaffolds and supported by the gene-prediction pipelines were retained.

### Mitochondrial genome assembly and annotation

Mitochondrial genome sequences and proteins from Simuliidae were downloaded from NCBI (Supplementary Table 1) and used as BLASTN and tBLASTN queries to identify mitochondrial sequence in the *S. vittatum* assembly. A contig with matches spanning reference mitogenomes and with the expected mitochondrial-genome length was extracted and annotated with MITOS2 (68) through the Galaxy implementation of MITOS (69). Orthologous genes were aligned with MAFFT (70), converted to PHYLIP format with Biopython (71), and used to infer a multigene phylogeny with 100 bootstrap replicates.

## Results

### Genome *de novo* assembly and quality assessment

The final curated *S. vittatum* assembly spans 340.4 Mb across 94 scaffolds and 1,654 contigs (Supplementary Table 2 and Supplementary Figure 1). Three chromosome-length scaffolds account for 99.1% of the total assembly, consistent with the three chromosome pairs (2n = 6) described cytogenetically for *S. vittatum* (12,20). The chromosome assembly is highly contiguous (N50 is 104.8 Mb), with only two gaps totaling 200 ambiguous “N” bases, with an overall GC content of 34.0% (Table 3). Manual curation of Hi-C contact map resolved initial structural inconsistencies through targeted scaffold breaking, reorientation, and rejoining, producing robust intrachromosomal interaction signals with minimal artifacts (Figure 1B). Benchmarking with BUSCO recovered 93.9% of expected dipteran orthologs as complete (92.9% as single copy and 1.0% duplicated), with 1.2% fragmented and 4.9% missing.

**Figure 1:**
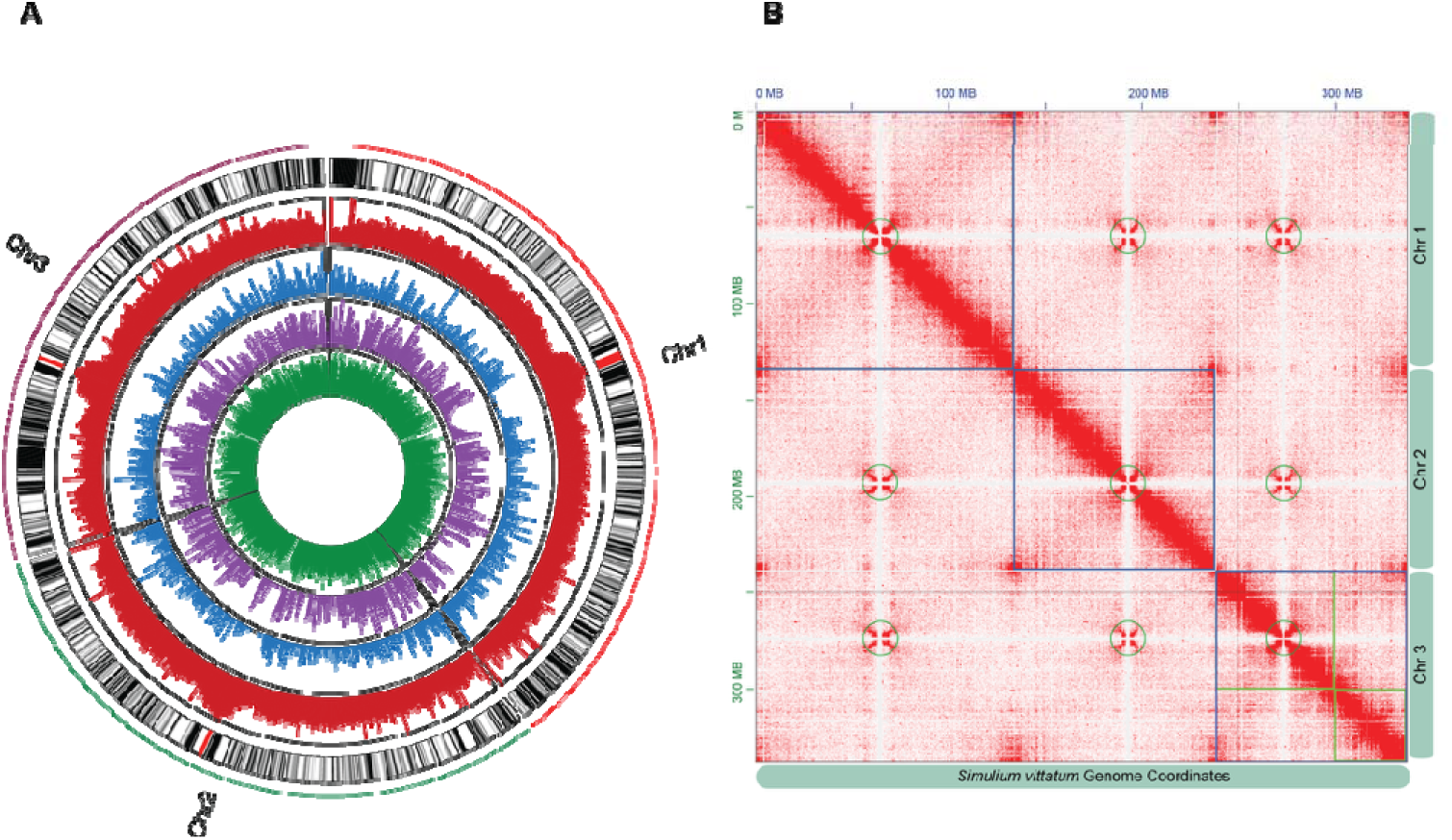
**A)** Circos plot showing the three chromosomes of Simulium vittatum, in this chromosome-level genome assembly. The outer ring shows the GC content in 250kbp windows (25-40%, from white to black), with the centromeres labelled in red. The second ring in red shows repeat density in 250kbp windows (23-100%). The third blue ring shows the coding density in 250kbp windows (0-30%). The fourth purple ring shows the SNP density per 250kbp windows (0-1363). The inner green ring shows the mean TPM of genes across the genome, averaging the expression of all RNA sequencing samples used for functional annotation. **B**) Hi-C contact heatmap of Simulium vittatum at 25 kb resolution showing three chromosome-scale scaffolds, consistent with the known 2n = 6 karyotype of blackflies. The x- and y-axes represent genomic coordinates. Green circles highlight centromeric interaction patterns within each chromosome. Color intensity reflects contact frequency.

**Table 3:**
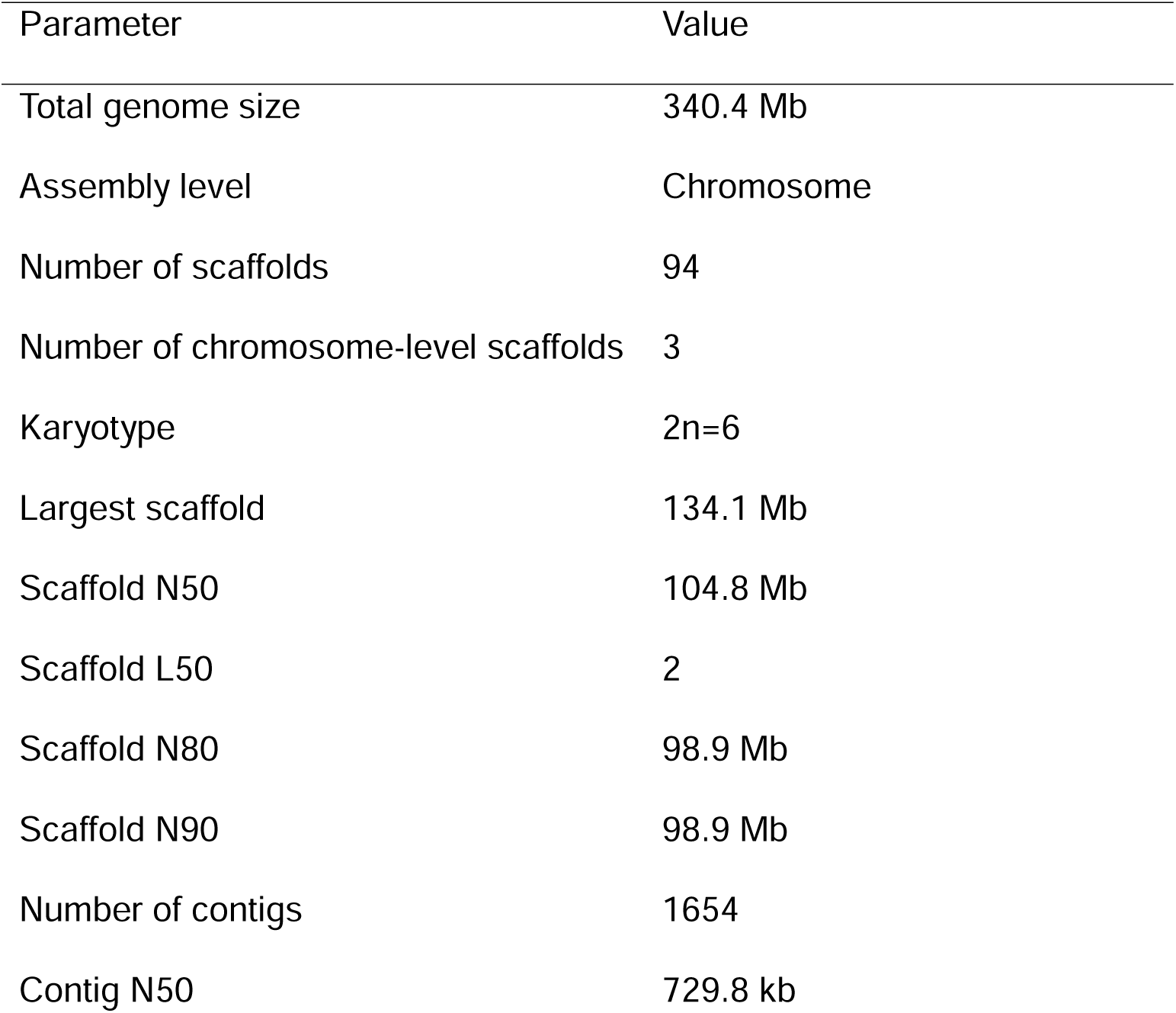
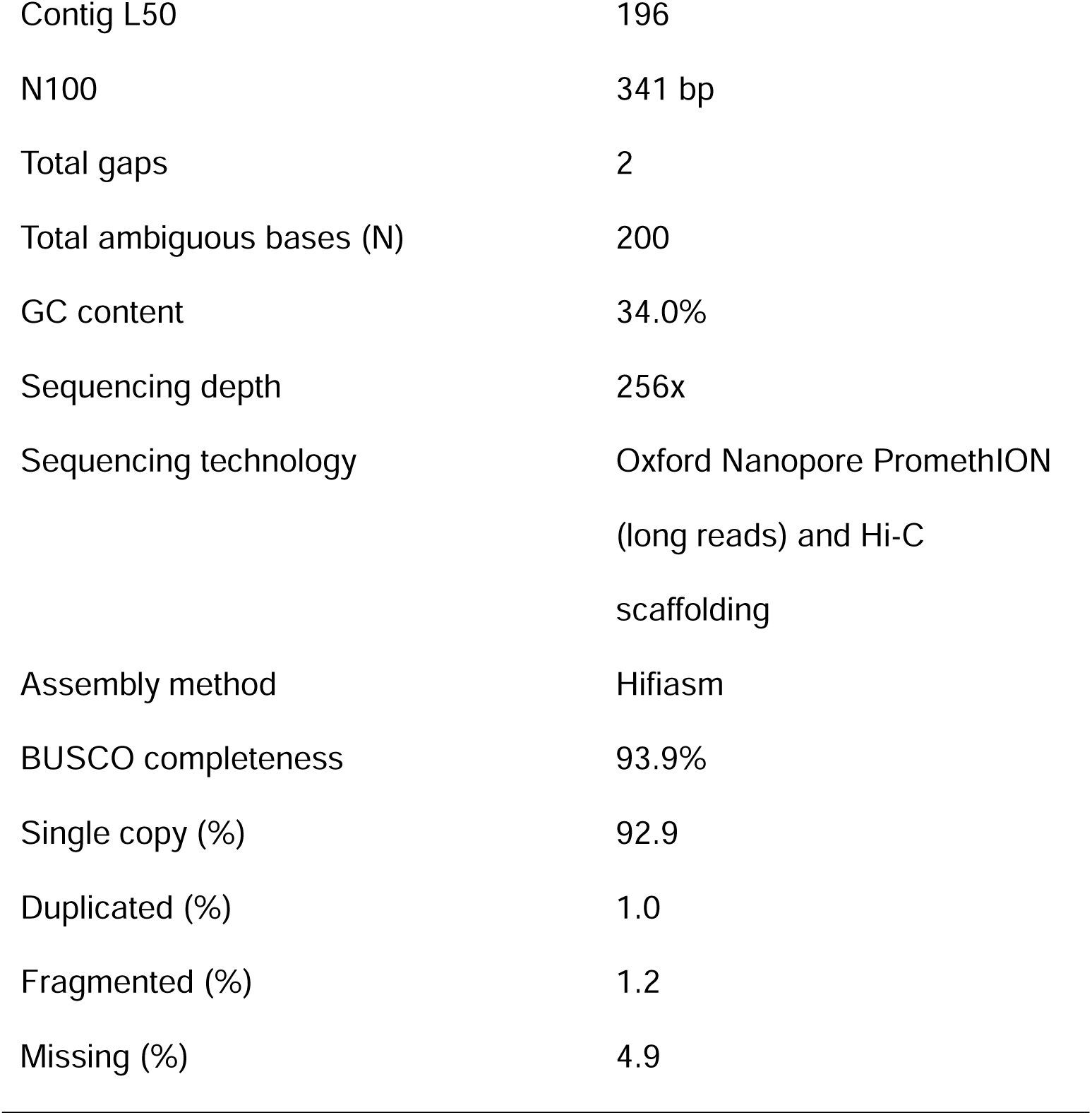
Summary metrics of the Simulium vittatum genome.

### Chromosome organization and cytogenetic correspondence

Chromosome numbering and arm assignments were based on chromosome sizes and inferred centromere locations. Chromosome I is approximately 134.1 Mb and metacentric, chromosome II is approximately 105 Mb and metacentric, and chromosome III is approximately 98.9 Mb and submetacentric. Hi-C contact patterns and changes in sequence composition supported candidate centromeric intervals on the three chromosome-scale scaffolds (Chr I: 64.25-66.00 Mb; Chr II: 58.25-59.00 Mb; Chr III: 34.50-35.25 Mb; Figure 1A and B). Repetitive elements were broadly distributed across all three chromosomes but showed increased density within these putative centromeric intervals, while canonical telomeric sequences were not recovered at the chromosome termini.

Molecular landmarks on chromosome III provide direct correspondence between the chromosome-level assembly and the classical polytene chromosome map. The predicted ortholog of the salivary antithrombin gene SVAT, historically mapped to band IIIS-72a4.5, localized to approximately 11.24 Mb in the assembly, whereas the predicted SVEP ortholog, mapped to band IIIL-96b1, lies at approximately 85.31 Mb (12). Flanking the putative centromeric region (∼34.5-35.25 Mb), these molecular landmarks unequivocally orient the low-coordinate scaffold arm as IIIS and the high-coordinate arm as IIIL.

### Genome annotation and functional gene repertoire

The integrated annotation identified 14,732 protein-coding genes and 16,417 transcripts, with 12,571 genes independently supported by both prediction pipelines (Supplementary Tables S3 and S4). Among the predicted genes, 14,652 had homologs in public protein databases, and 14,700 protein-coding loci were anchored to the three chromosome-scale scaffolds. Major functional annotation classes are summarized in Figure 2, and representative gene families are listed in Supplementary Table S5.

**Figure 2:**
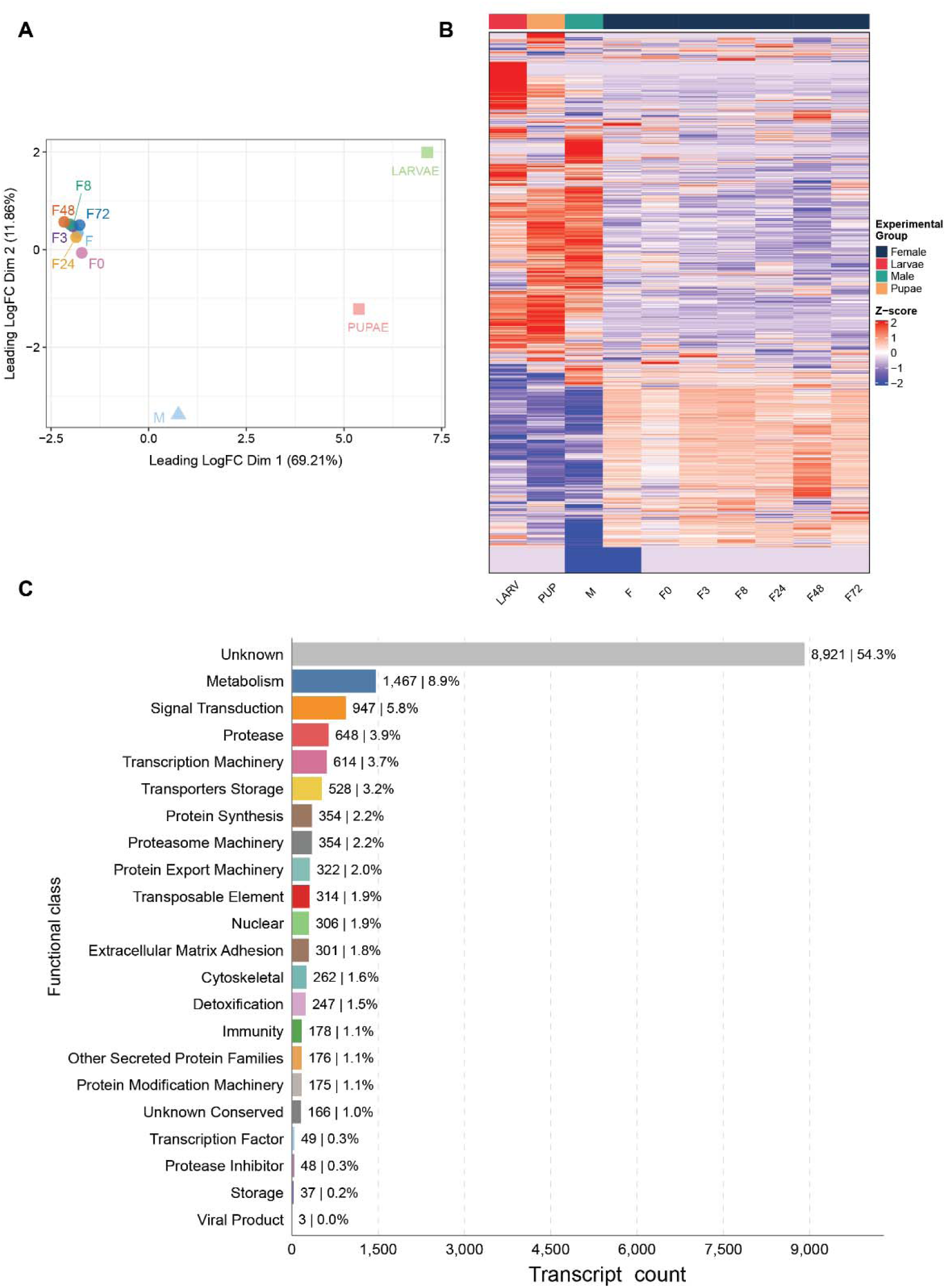
A) Multidimensional scaling (MDS) plot of S. vittatum RNA-seq samples representing larvae, pupae, adult males, adult females, and blood-fed females sampled at 0, 3, 8, 24, 48, and 72 h post-feeding. Numbers after f indicate hours after blood feeding. Because a single biological replicate was available for each condition, the MDS is presented descriptively and was not used for statistical inference. B) Heatmap of expressed genes across sampled conditions. Expression values are shown as row-wise z-scores from the available replicate for each condition; samples are ordered according to the lifecycle and blood-feeding progression shown in the panel. C) Distribution of predicted transcripts across broad functional classes. Bars show transcript counts and percentages. Functional classification was performed independently of predicted localization; therefore, secretion-associated annotations did not override more specific functional assignments.

The annotation recovered several gene families associated with major aspects of black fly biology. Hematophagy-related families included serine proteases, trypsins, chymotrypsins, cathepsins, apyrases, D7-family proteins, and other salivary proteins (11,72–74), several of which contained predicted N-terminal signal peptides. Genes associated with xenobiotic metabolism included cytochrome P450 monooxygenases, glutathione S-transferases, carboxylesterases, and ATP-binding cassette transporters (75). Chemosensory repertoires include odorant and gustatory receptors and opsins (76). Development-related families included cuticular proteins, chitin-metabolic enzymes, and components of ecdysone and juvenile-hormone signaling (77), whereas the innate immune repertoire includes CLIP-domain serine proteases, lysozymes, defensins, and cecropins (78–80).

Of the 2,788 transcripts independently assigned secretion-related functional annotations (Supplementary Tables S3 and S4), 2,541 also contained a SignalP-predicted signal peptide, defining a conservative set supported by both approaches. The remaining 247 transcripts retained secretion-related annotations but lacked a predicted classical signal peptide. Functional classes were assigned independently of predicted cellular localization. Overall, 54.3% of predicted transcripts remained functionally unclassified, under the parameters applied in this study, indicating that a substantial proportion of the *S. vittatum* proteome remains to be functionally characterized.

The chromosome-level annotation recovered the three previously characterized *S. vittatum* larval silk gland proteins (81) (Supplementary Note S1). SGP-1, SGP-2, and SGP-3 corresponded to SV00009995-RA, SV00009996-RA, and SV00009306-RA, respectively, and all three encoded proteins with predicted N-terminal signal peptides. SGP-1 and SGP-3 showed 97.9% and 100% amino-acid identity, respectively, to the published proteins, whereas an RNA-supported reconstruction of SGP-2 showed 97.0% identity and 98.2% similarity to the published SGP-2 sequence (Supplementary Alignments S1–S3). All three transcripts showed markedly greater abundance in the larval sample than in pupal and adult samples (Supplementary Table S3), in agreement with their established roles in silk production. SGP-1 and SGP-2 were separated by only 3,054 bp on chromosome II, whereas SGP-3 occurred approximately 13.5 Mb away on the same chromosome (Supplementary Figure S2).

Detailed evaluation of SGP-2 revealed a sequence discrepancy between the genome-derived model and the expressed transcript. The initial genome-derived SV00009996-RA model predicted a 365-aa protein that closely matched published SGP-2 through residue 128 but diverged thereafter, coincident with the junction between two annotated coding exons.

Incorporation of an additional G relative to the genome-derived sequence at this junction restored a 336-aa open reading frame with 97.0% identity and 98.2% similarity to published SGP-2. Notably, the previously deposited SGP-2 mRNA sequence (KC699730.1) contains the same additional G at this position, providing independent sequence support for the reconstructed coding frame.

Larval RNA-seq reads further supported the RNA-associated junction sequence, whereas no exact matches to the corresponding genome-derived junction sequence were detected.

Representative 150-bp junction-spanning RNA reads mapped with 100% identity to the SGP-2 genomic regions flanking the annotated intron. Furthermore, genomic alignments between the raw ONT contigs and final Hi-C scaffolds were identical across this region, indicating that the discrepancy was not introduced during scaffolding (Supplementary Note S1; Supplementary Figure S2).

While the RNA-seq read mapping supported an alternative 336-amino-acid SGP-2 coding sequence, the discrepancy between the genomic assembly and the transcript-supported sequence at one nucleotide position remains unresolved.

### Structural variation and sex-associated genomic features

Pairwise alignment between the primary (Hap1) and alternate (Hap2) assembled haplotypes revealed no large heterozygous inversion corresponding to the historical chromosome-I IS-7 rearrangement. Because cross-haplotype comparison detects structural differences between homologous chromosomes, the absence of such a rearrangement is compatible with homozygosity across the historical IS-7 interval in the sequenced male. Cytogenetically, the individual could therefore represent either the standard (SS) or inverted (II) arrangement on both homologs. However, historical cytogenetic data documented SS and SI males but no II males (20), making SS the more plausible interpretation. Definitive assignment will require mapping the historical cytogenetic inversion breakpoints to sequence coordinates.

Genome-wide structural comparison of the two male haplotypes identified 71 putative heterozygous inversions ranging from approximately 0.1 to 10 Mb. A ∼9.3 Mb inversion spans approximately 14.6-23.9 Mb on the short arm of chromosome III (Figure 3). Because SVAT and SVEP anchor this arm as IIIS, this rearrangement cannot be equated with the historical IIIL-1 sex-linked system. Its position on IIIS and proximity to SVAT instead make the polymorphic IIIS-2 inversion a plausible cytogenetic comparator (12), although direct correspondence between the two rearrangements has not been demonstrated (Figure 3).

**Figure 3.**
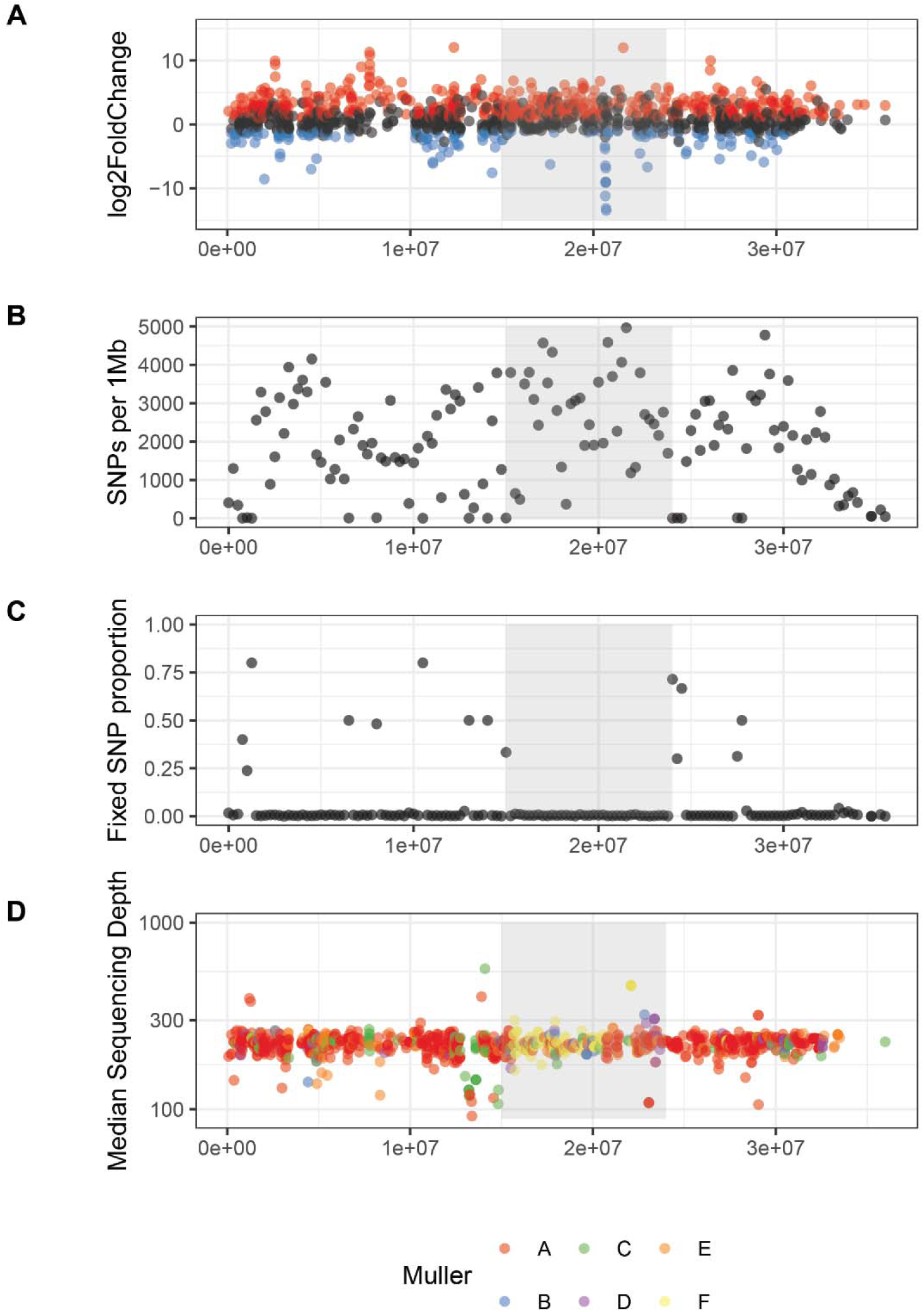
Candidate sex-associated region localized on chromosome IIIS in Simulium vittatum. **A)** The plot shows genes with higher transcript abundance towards males (blue) or female (red). **B)** SNPs distribution per 1Mb window. **C)** Proportion of SNPs in each 1Mb window that are fixed (e.g. not heterozygous). **D)** Genomic sequencing coverage of genes and the conserved Muller element that gene is found on in Drosophila melanogaster and Aedes aegypti. The shaded window across all plots shows the breakpoints for the heterozygous inversion detected by Delly.

**Figure 4:**
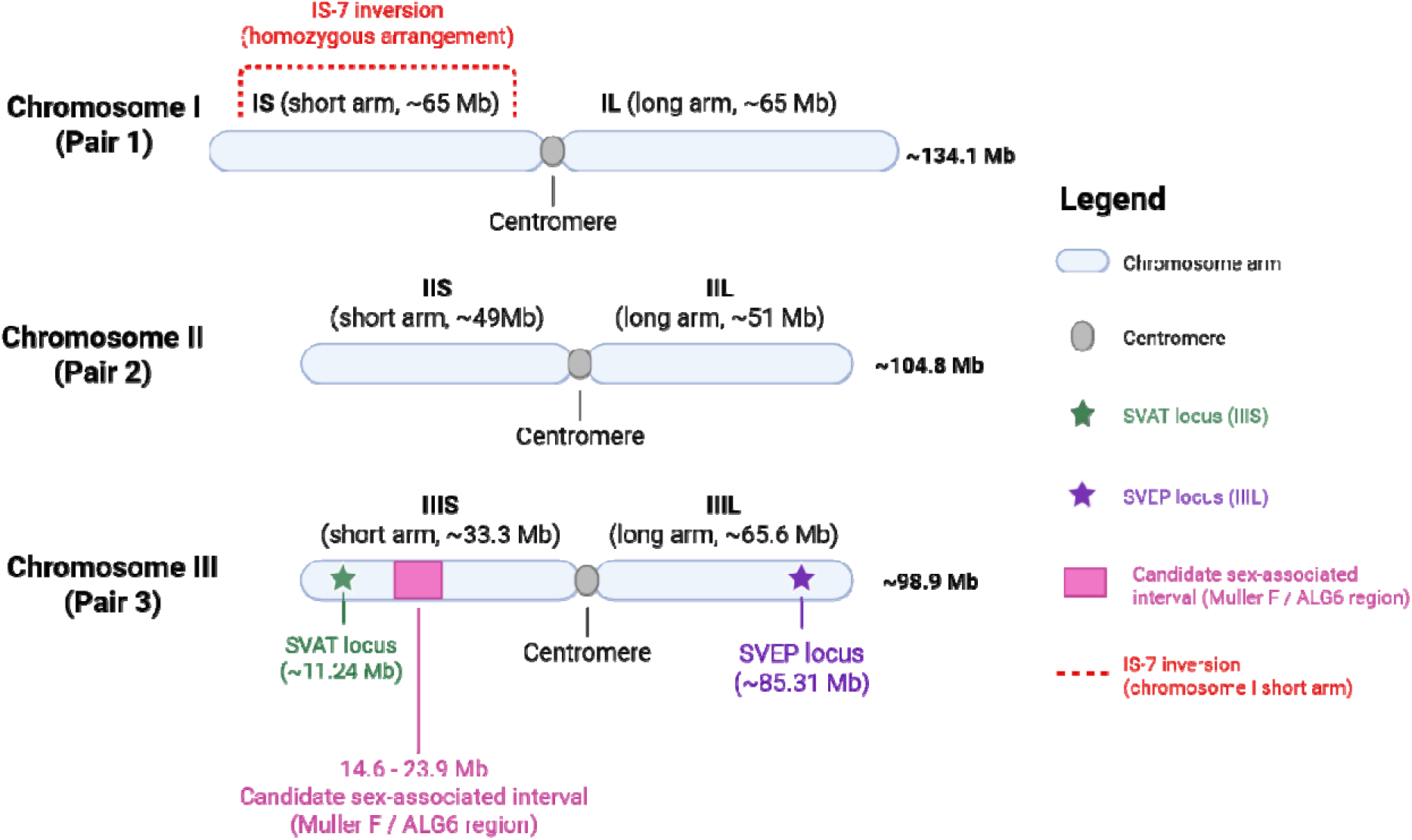
Chromosomal organization and cytogenetic landmarks of the S. vittatum IS-7 genome. Schemati representation of the three chromosome-length scaffolds corresponding to the haploid chromosome complement of S. vittatum (chromosomes I-III). Approximate chromosome and arm lengths are shown according to established cytogenetic nomenclature. On chromosome III, the previously mapped SVAT and SVEP loci provide molecular anchors for the short (IIIS) and long (IIIL) arms, respectively, linking the chromosome-level assembly to the established cytogenetic framework. SVAT maps to approximately 11.24 Mb on IIIS, whereas SVEP maps t approximately 85.31 Mb on IIIL. The candidate sex-associated interval identified by Muller-element analysis spans approximately 14.6-23.9 Mb on IIIS. The historical IS-7 inversion is indicated schematically on the short arm of chromosome I; its absence as a heterozygous rearrangement in the sequenced male is consistent with a homozygous arrangement at this locus. Chromosome and arm lengths are approximate, and the schematic is not intended to represent cytological banding patterns or the physical dimensions of individual chromosome features. Created in BioRender. Sayuri Nishiduka Costa, E. (2026) https://BioRender.com/zttzuje.

Descriptive comparison of adult male and female transcript abundance identified a cluster of genes with higher transcript abundance in males within part of the IIIS interval (Figure 3A). This region also showed increased heterozygous SNP density (Figure 3B and C) and contained multiple orthologs of genes located on Drosophila Muller element F (Figure 3D). Muller F represents a derived component of ancestral linkage group 6 (ALG6), which has been implicated in dipteran sex-chromosome evolution and sex determination (82).

### Repetitive DNA and transposable elements

*De novo* repeat discovery initially identified approximately 1,270 candidate repeat consensus sequences. After filtering and redundancy reduction, the curated nonredundant library contained 711 consensus sequences, comprising 276 classified TE consensus sequences and 435 unclassified repeat consensus sequences. Manual refinement yielded 80 putatively full-length TE consensus models, comprising four Class II elements, 42 non-LTR retrotransposons, and 34 LTR retrotransposons. All putatively full-length models were supported by near-complete genomic copies spanning at least 90% of the corresponding consensus sequence, and most were represented by multiple such copies. Class-specific structural features and independently validated coding regions provided additional support for the reconstructed element boundaries. Among these models, we recovered a 15.2-kb Polinton/Maverick consensus that was supported by multiple near-complete genomic copies.

Genome-wide repeat annotation showed that 46.48% of the *S. vittatum* assembly consisted of repetitive or low-complexity sequence (Figure 5A). Classified TEs accounted for 15.39% of the genome, with DNA transposons representing the largest component: TIR elements occupied 8.48% of the assembly and Mavericks an additional 0.57%, together comprising 58.8% of the sequence assigned to classified TEs. Genomic abundance, however, did not mirror the number of curated consensus models. LTR retrotransposons comprised the largest number of models (107) but occupied only 2.40% of the assembly, compared with 8.48% occupied by 74 TIR models and 3.77% by 83 LINE models. Within the TIR fraction, the main contributors were TIR-bearing elements without confident superfamily assignment, together with MITEs and hAT-related elements, whereas CR1, RTE, Kiri, and LOA dominated the LINE fraction. Gypsy and Bel-Pao together accounted for most of the LTR-derived sequence. PLEs (0.14%) and SINEs (0.02%) were minor components, while simple repeats and low-complexity regions contributed an additional 1.41% of the assembly (Supplementary Table S6). Unclassified repeats accounted for 29.68% of the assembly and comprised 435 curated consensus sequences with no detectable coding regions longer than 600 nt, recognizable protein domains in CDD or Pfam, diagnostic TE structures, or informative Swiss-Prot matches. They were therefore conservatively retained as unclassified repeats, although some may represent highly diverged or lineage-specific TE remnants.

**Figure 5:**
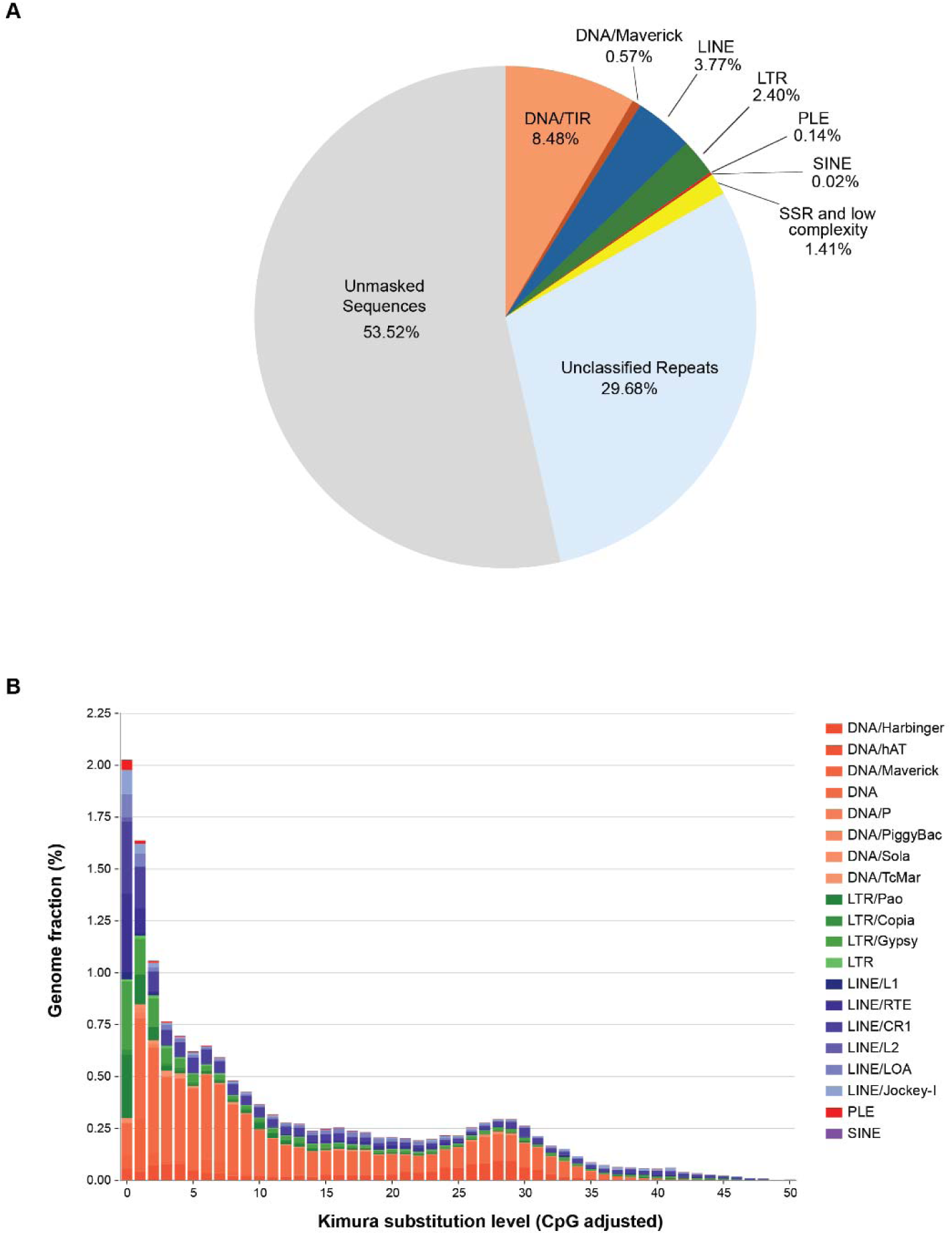
Repeat composition and transposable element divergence landscape of the Simulium vittatum genome. **A)** Genome composition based on RepeatMasker annotation using the final curated repeat library. Percentages indicate the fraction of the genome occupied by terminal inverted repeat (TIR) DNA transposons, Mavericks, long interspersed nuclear elements (LINEs), long terminal repeat (LTR) retrotransposons, Penelope-like elements (PLEs), short interspersed nuclear elements (SINEs), simple sequence repeats (SSRs) and low-complexity sequences, unclassified repeats, and unmasked sequences. **B)** Divergence landscape of classified transposable elements based on RepeatMasker alignments of genomic copies to their corresponding curated consensus sequences. Stacked bars show the percentage of the genome occupied by individual TE superfamilies across CpG-adjusted Kimura divergence classes. Lower divergence from the consensus is indicative of relatively more recent TE insertions, whereas higher divergence is consistent with older insertions. Colors denote TE classes and superfamilies as indicated in the key.

Analysis of Kimura 2-parameter (K2P) divergence between genomic TE copies and their consensus sequences revealed a strong enrichment of TE-derived sequence at low divergence, with 30.4% occurring within the 0-2% interval (Figure 5B). This pattern was particularly pronounced among retrotransposons, with 49.9% of LTR-derived and 42.3% of LINE-derived sequence falling within this interval, compared with 20.0% of DNA transposon sequence.

Several lineages contributed substantially to the low-divergence fraction, particularly MITEs, Gypsy- and Bel-Pao-like LTR retrotransposons, RTE LINEs, and Mavericks. Some of these lineages also included putatively full-length models represented by multiple near-complete genomic copies, consistent with comparatively recent amplification and retention of structurally preserved elements. In contrast, a second increase at approximately 25-30% divergence was dominated by DNA transposons, particularly TIR-bearing elements without confident superfamily assignment and hAT-related elements. Together, these patterns indicate distinct divergence profiles among TE lineages and are compatible with differences in their amplification histories within the *S. vittatum* genome.

### Candidate horizontal gene transfers

Our analysis identified nine candidate horizontally transferred proteins that met both the AI > 30 and hU > 30 thresholds (Supplementary Table S7). Most candidate proteins showed their strongest taxonomically informative similarities to bacterial homologs, particularly proteins from *Wolbachia* and other Rickettsiales. Several of those encode ankyrin-repeat or tetratricopeptide-repeat proteins, domains frequently involved in protein–protein interactions at host-symbiont interfaces (83,84). A smaller number of candidates match fungal- or viral-like proteins.

Five of the candidate HGT genes occur within a contiguous gene-model block, SV00006927– SV00006932, embedded in a larger tandem array of ankyrin-repeat genes. Several candidates have best outgroup matches to *Wolbachia* ankyrin- or tetratricopeptide-repeat proteins, whereas the broader functional annotation also identifies insect homologs for members of this locus. A retrotransposon-related gene (SV00006933) occurs immediately adjacent to the array. This organization raises the possibility that the HGT signals reflect either a historical transfer of a larger symbiont-derived genomic fragment followed by duplication and divergence or, alternatively, phylogenetic ambiguity associated with an expanded host ankyrin-repeat family.

The clustered organization therefore supports treating these candidates as a single genomic locus rather than as independent HGT events.

### Mitochondrial genome and phylogenetic placement

The *S. vittatum* mitochondrial genome is 15,842 bp long and contains the expected 37 metazoan mitochondrial genes: 13 protein-coding genes, 22 tRNAs, and two ribosomal RNA genes (Figure 6A; Supplementary Table S8). The genome contains six short gene overlaps and 29 predominantly short intergenic spacers. Twenty-three genes are encoded on the majority strand and 14 on the minority strand. No major gene-order rearrangements were detected relative to the sampled Simuliidae mitochondrial genomes and the phylogenetic analysis placed *S. vittatum* within the expected Simuliidae lineage (Figure 6B).

**Figure 6:**
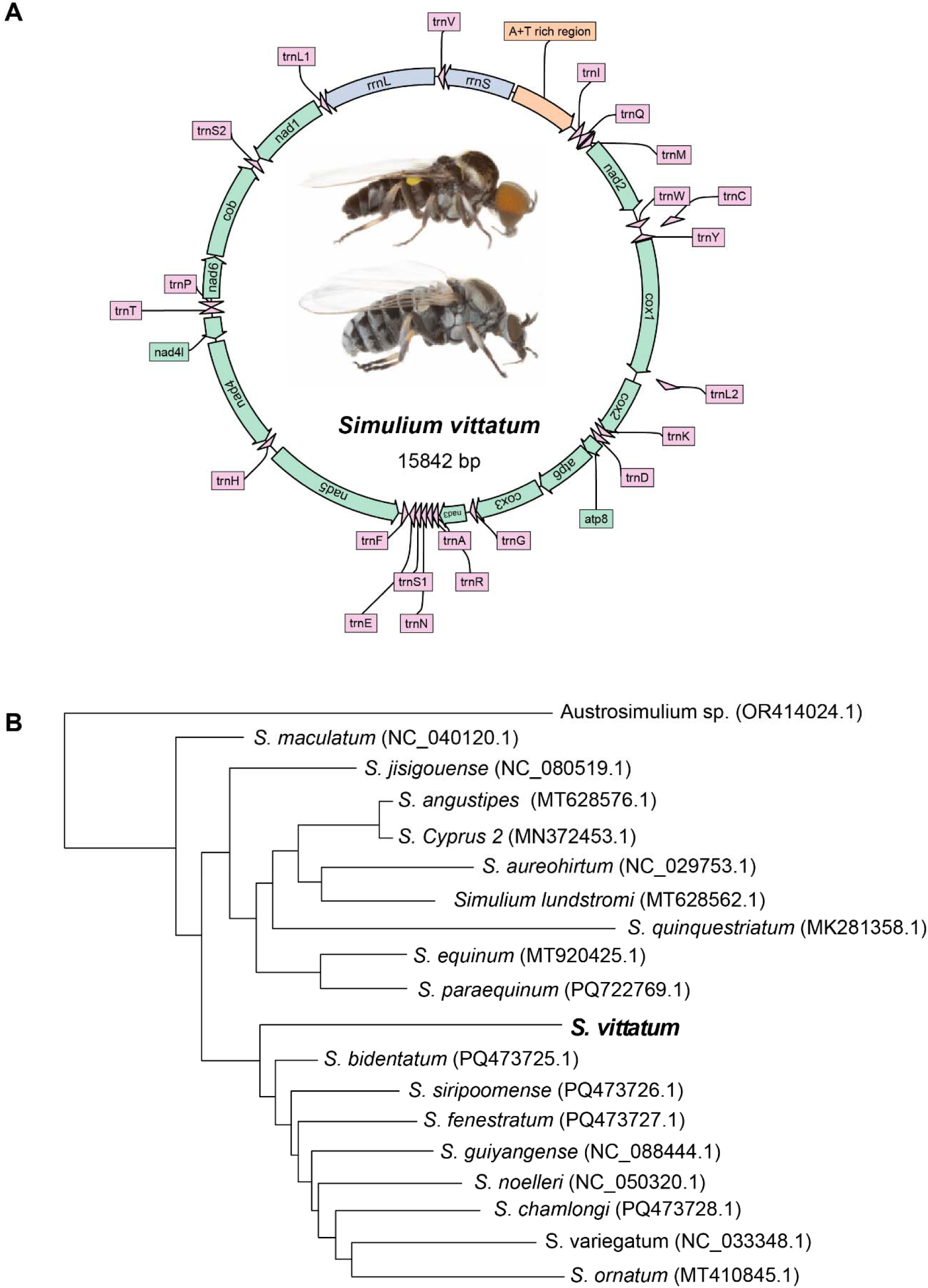
Mitochondrial genome characteristics of Simulium vittatum. The circular mitochondrial genome has a total length of 15,842 bp. Protein-coding genes are represented in green, transfer RNA in pink, and ribosomal RNA genes in purle. **(a)** Mitochondrial genome map showing annotated genes and their transcriptional orientation. **(b)** Phylogenetic analysis of selected Simulium species based on mitochondrial sequences, with NCBI GenBank accession numbers shown in parentheses.

## Discussion

The principal value of this chromosome-level assembly is that it connects a high-resolution sequence resource with the well-developed cytogenetic traditions of Simuliidae. More than 99% of the 340.4-Mb assembly is contained in three chromosome-length scaffolds whose sizes and centromere positioning agree with the 2n = 6 karyotype described for *S. vittatum*. The sequence positions of SVAT and SVEP provide independent landmarks for chromosome III and identify the low- and high-coordinate arms as IIIS and IIIL, respectively. This correspondence is important because it allows structural variants detected in the assembly to be discussed in the same nomenclatural framework as historical polytene chromosome studies, directly linking sequence-level genome organization with the extensive cytogenetic knowledge available for black flies.

The chromosome-I result is also a useful example of why structural variation from a single diploid individual must be interpreted cautiously. Failure to detect the historical IS-7 inversion as a heterozygous rearrangement does not imply that the sex-linked system has been lost from the colony. Rather, the absence of a heterozygous rearrangement is compatible with the sequenced male being homozygous for the arrangement present at this locus. Because historical samples contained SS and SI males but no II males (20), an SS genotype is the more likely explanation. Direct mapping of the historical inversion breakpoints and genotyping of additional males and females are needed to test this interpretation at sequence resolution.

The 14.6-23.9-Mb heterozygous inversion on IIIS should therefore be considered separately from the chromosome-I IS-7 system. Its location and proximity to SVAT make the common IIIS-2 autosomal inversion a plausible historical counterpart (12), but the current data do not establish equivalence. Nor do they establish the IIIS interval as a sex-determining region. The cluster of transcripts with higher abundance in the male sample is descriptive, and the presence of Muller F-homologous genes indicates evolutionary homology rather than sex linkage. Muller F is a derived subset of ALG6, and broad dipteran comparisons suggest that ALG6 was ancestrally associated with sex chromosomes before extensive lineage-specific fusion and redistribution (82). Because Simuliidae were not included in that reconstruction, mapping ALG1-ALG6 markers across the *S. vittatum* genome will be important for placing these observations in a broader evolutionary context.

The repetitive landscape of *S. vittatum* genome is also notable. Repetitive and low-complexity sequences comprise 46.48% of the assembly, a larger fraction than reported for several closer Culicomorpha comparators, including *C. brevitarsis* (approximately 14.6%), *C. riparius* (approximately 15.85%), and *C. sonorensis*, for which RepeatModeler identified approximately 15% repetitive sequence and a broader repeat-plus-low-complexity estimate reached 29.7% (21,23). Mosquitoes span a wider range, with substantially less repetitive sequence in *A. gambiae* than in the TE-rich *Ae. aegypti* genome (24,25). These cross-species comparisons are necessarily approximate because repeat estimates depend strongly on assembly quality, repeat libraries, and annotation pipelines, and the extent of manual curation. Nevertheless, *S. vittatum* has a relatively large repetitive fraction compared with these available Culicomorpha references. Importantly, this repetitive fraction is not equivalent to classified TE content: only 15.39% of the assembly was assigned to established TE groups, whereas 29.68% consisted of unclassified repeats. Among classified TEs, DNA transposons accounted for the largest genomic fraction.

Thus, the relatively large size of the *S. vittatum* genome may reflect broad accumulation of repetitive DNA rather than expansion of a single recognized TE class.

The divergence profiles suggest that the *S. vittatum* repeatome reflects multiple histories of TE amplification. Highly diverged DNA transposon-derived sequences coexist with much less divergent retrotransposon lineages, indicating that different TE groups contributed to genome evolution at different times. The presence putative full-length models among some of the least divergent lineages further suggests that some recently amplified copies have retained relatively intact structural features. High proportions of unclassified repeats are common in non-model insects and have been linked to uneven taxonomic representation in repeat databases, limiting recognition of lineage-specific or highly diverged sequences (85). In *S. vittatum*, the limited availability of closely related genomic references may further hinder repeat classification. Because these sequences also lack recognizable coding, structural, and protein-homology signatures, at least some may represent ancient or lineage-specific TE-derived fragments whose diagnostic features have eroded beyond current detection.

The HGT screen addresses a second potential source of genome novelty, although the evidence is substantially more preliminary than that supporting the TE landscape, as the HGT analysis identified candidates rather than confirmed integration events. The predominance of matches to *Wolbachia* and related intracellular bacteria is biologically plausible because natural *Wolbachia* infections occur in black fly populations (86), and endosymbiont-to-host transfer has been documented in other insects. Residual symbiont DNA, contamination, or annotation artifacts remain alternative explanations for the identified HGT candidates. Confirmation will require host-derived flanking sequence and continuous long-read support across putative integration boundaries, followed where appropriate by independent molecular validation.

The mitochondrial genome provides a complementary evolutionary record. Its conserved organization contrasts with the extensive inversion polymorphism of simuliid nuclear chromosomes, and previous work in *S. vittatum* showed that mitochondrial variation does not necessarily track cytologically defined sibling groups (30). Nuclear and mitochondrial references from the same experimental lineage now allow these historically separate sources of information to be compared directly.

The protein-coding annotation also places decades of physiological work on IS-7 into a genomic context. Salivary and digestive proteins associated with hematophagy, chemosensory receptors, detoxification enzymes, developmental genes, and innate immune factors can now be examined as defined chromosomal loci rather than as isolated gene products or transcripts. The recovery of the previously characterized larval silk proteins SGP-1-SGP-3 provides a direct example of this utility. All three were assigned to chromosome II, with SGP-1 and SGP-2 separated by only 3,054 bp, revealing genomic organization that was not apparent from the previously characterized protein sequences alone. Their greater transcript abundance in the larval sample is consistent with their established roles in silk production. Examination of SGP-2 further identified a single-nucleotide discrepancy between the genome-derived gene model and the expressed transcript at a coding-exon junction. Larval RNA-seq supported a reconstructed 336- aa coding sequence that closely matches the previously characterized SGP-2 protein, illustrating how the combined genome and developmental transcriptome can be used not only to place previously characterized genes in chromosomal context but also to reassess their sequence-level annotation. More broadly, the developmental RNA-seq data extend annotation support across life stages and blood-feeding conditions and provide a resource for prioritizing genes for future functional studies.

Several limitations define the scope of the present study. The nuclear genome was sequenced from a single male, so sex association cannot be inferred from structural heterozygosity in that individual alone. The RNA-seq design included one pooled biological sample per stage or condition and therefore supports annotation and descriptive expression profiling, but not formal differential-expression inference. The historical IS-7 and IIIS-2 inversion breakpoints have not yet been mapped to sequence coordinates; the genomic basis of the SGP-2 genome-transcript discrepancy remains unresolved; and the HGT candidates have not been validated experimentally. Cross-species repeat comparisons were based on independently curated repeat annotations rather than a single standardized annotation pipeline, which limits strictly quantitative comparisons among species. However, species-specific curation was considered more appropriate for these non-model taxa because automated repeat annotation can generate substantial numbers of false-positive predictions that require manual evaluation. These limitations primarily constrain the biological interpretation of particular loci and candidate regions rather than the overall contiguity of the reference assembly.

## Conclusions

To our knowledge, this study provides the first chromosome-scale nuclear genome assembly for a member of the Simuliidae and connects the assembly directly to the extensive cytogenetic framework available for *S. vittatum*. The integration of long-read sequencing, Hi-C scaffolding, gene and repeat annotation, and life-stage and post-blood-feeding transcriptomes provides a genomic framework for investigating chromosome evolution, repetitive-DNA dynamics, candidate horizontal gene transfer, salivary biology, and sex-associated variation in black flies. This resource strengthens the value of the IS-7 colony as an experimental system for vector biology and comparative dipteran genomics. Furthermore, the resulting reference genome addresses a major genomic gap within Culicomorpha and provides a foundation for comparative and population genomic studies of chromosome evolution and other vector-associated traits across Simuliidae.

## List of abbreviations

ABC: ATP-binding cassette
AI: Alien Index
BUSCO: Benchmarking Universal Single-Copy Orthologs
FPKM: fragments per kilobase per million reads
HGT: horizontal gene transfer
HMW: high-molecular-weight
ONT: Oxford Nanopore Technologies
PCG: protein-coding gene
SNP: single-nucleotide polymorphism
TE: transposable element
TIR: terminal inverted repeat
TPM: transcripts per million
VSV: vesicular stomatitis virus.

## Data Availability

Supplementary Data are available at NAR Online. All read data and final assemblies have been deposited in the National Center for Biotechnology Information (NCBI) under BioProject (PRJNA1439345). The raw sequencing dataset generated for this project, including Illumina and Oxford Nanopore (ONT) data, was submitted to the NCBI Sequence Read Archive (https://identifiers.org/insdc.sra) under the following accession numbers: ONT reads (SRR37678976), Hi-C reads (SRR37678975) used to scaffold the assemblies; and Illumina short-reads for the developmental stage dataset: (SRR37678957-SRR37678974).

Finally, custom R scripts used in horizontal gene transfer analysis are available at: https://github.com/liliancaesarbio/alien_hgt_index.

## Acknowledgements

This research was supported [in part] by the Intramural Research Program of the National Institutes of Health (NIH). The contributions of the NIH author(s) are considered Works of the United States Government. The findings and conclusions presented in this paper are those of the author(s) and do not necessarily reflect the views of the NIH or the U.S. Department of Health and Human Services. The *Simulium vittatum* colony used in this work was maintained with the support of NIH Task Order C-08, Contract [No. HHSN2722017000351], Task Order [No. 75N93020F00002] and obtained through BEI Resources, NIAID, NIH: *Simulium vittatum*, Adult, NR-53893. This work utilizes the computational resources of the NIH HPC Biowulf cluster (http://hpc.nih.gov).

## Author contributions

Conceptualization: E.N, E.C.

Methodology: E.N, E.C, O.M, T.H, S.L, B.B, E. M, L.C., L.T, G.L.W, P.V

Investigation: E.N, E.C, O.M, T.H, S.L, E. M, G.L.W, L.C.

Writing – original draft: E.N, E.C, O.M, T.H, E.M, L.C.

Writing – review and editing: E.N, E.C, O.M, T.H, E.M, S.L., B.B, L.C., L.T, G.L.W, P.V

Funding acquisition: E.C.

Resources: E.C.

## References

1. Adler, P.H. and McCreadie, J.W. (2019) In Mullen, G. R. and Durden, L. A. (eds.), Medical and Veterinary Entomology (Third Edition). Academic Press, pp. 237–259.

2. Adler, P.H., Cheke, R.A. and Post, R.J. (2010) Evolution, epidemiology, and population genetics of black flies (Diptera: Simuliidae). Infect. Genet. Evol., 10, 846–865.

3. Cupp, E.W. and Cupp, M.S. (1997) Black fly (Diptera:Simuliidae) salivary secretions: importance in vector competence and disease. J. Med. Entomol., 34, 87–94.

4. Enk, C.D. (2006) Onchocerciasis--river blindness. Clin. Dermatol., 24, 176–180.

5. Hellgren, O., Bensch, S. and Malmqvist, B. (2008) Bird hosts, blood parasites and their vectors--associations uncovered by molecular analyses of blackfly blood meals. Mol. Ecol., 17, 1605–1613.

6. Cupp, E.W., Maré, C.J., Cupp, M.S. and Ramberg, F.B. (1992) Biological transmission of vesicular stomatitis virus (New Jersey) by Simulium vittatum (Diptera: Simuliidae). J Med Entomol, 29, 137–140.

7. Murdock, C.C., Adler, P.H., Frank, J. and Perkins, S.L. (2015) Molecular analyses on host-seeking black flies (Diptera: Simuliidae) reveal a diverse assemblage of Leucocytozoon (Apicomplexa: Haemospororida) parasites in an alpine ecosystem. Parasit Vectors, 8, 343.

8. Adler, P.H., Yildirim, A., Onder, Z., Tasci, G.T., Duzlu, O., Arslan, M.O., Ciloglu, A., Sari, B., Parmaksizoglu, N. and Inci, A. (2016) Rearrangement hotspots in the sex chromosome of the Palearctic black fly Simulium bergi (Diptera, Simuliidae). Comp Cytogenet, 10, 295–310.

9. Adler, P.H. and Huang, S. (2022) Chromosomes as Barcodes: Discovery of a New Species of Black Fly (Diptera: Simuliidae) from California, USA. Insects, 13.

10. Post, R.J. (1982) Sex-linked inversions in blackflies (Diptera:Simuliidae). Heredity (Edinb), 48, 85–93.

11. Andersen, J.F., Pham, V.M., Meng, Z., Champagne, D.E. and Ribeiro, J.M. (2009) Insight into the sialome of the Black Fly, Simulium vittatum. J. Proteome Res., 8, 1474–1488.

12. Procunier, W., Zhang, D., Cupp, M.S., Miller, M. and Cupp, E.W. (2005) Chromosomal localization of two antihemostatic salivary factors in Simulium vittatum (Diptera: Simuliidae). J Med Entomol, 42, 805–811.

13. Cupp, M.S., Ribeiro, J.M., Champagne, D.E. and Cupp, E.W. (1998) Analyses of cDNA and recombinant protein for a potent vasoactive protein in saliva of a blood-feeding black fly, Simulium vittatum. J. Exp. Biol., 201, 1553–1561.

14. Abebe, M., Ribeiro, J.M., Cupp, M.S. and Cupp, E.W. (1996) Novel anticoagulant from salivary glands of Simulium vittatum (Diptera: Simuliidae) inhibits activity of coagulation factor V. J. Med. Entomol., 33, 173–176.

15. Abebe, M., Cupp, M.S., Champagne, D. and Cupp, E.W. (1995) Simulidin: a black fly (Simulium vittatum) salivary gland protein with anti-thrombin activity. J. Insect Physiol., 41, 1001–1006.

16. Cupp, M.S., Ribeiro, J.M. and Cupp, E.W. (1994) Vasodilative activity in black fly salivary glands. Am. J. Trop. Med. Hyg., 50, 241–246.

17. Jacobs, J.W., Cupp, E.W., Sardana, M. and Friedman, P.A. (1990) Isolation and characterization of a coagulation factor Xa inhibitor from black fly salivary glands. Thromb. Haemost., 64, 235–238.

18. Ribeiro, J.M., Charlab, R., Rowton, E.D. and Cupp, E.W. (2000) Simulium vittatum (Diptera: Simuliidae) and Lutzomyia longipalpis (Diptera: Psychodidae) salivary gland hyaluronidase activity. J. Med. Entomol., 37, 743–747.

19. Bernardo, M.J., Cupp, E.W. and Kiszewski, A.E. (1986) Rearing black flies (Diptera: Simuliidae) in the laboratory: bionomics and life table statistics for Simulium pictipes. J Med Entomol, 23, 680–684.

20. Brockhouse, C.L. and Adler, P.H. (2002) Cytogenetics of laboratory colonies of simulium vittatum cytospecies IS-7 (Diptera: Simuliidae). J Med Entomol, 39, 293–297.

21. Morales-Hojas, R., Hinsley, M., Armean, I.M., Silk, R., Harrup, L.E., Gonzalez-Uriarte, A., Veronesi, E., Campbell, L., Nayduch, D., Saski, C. et al. (2018) The genome of the biting midge Culicoides sonorensis and gene expression analyses of vector competence for bluetongue virus. BMC Genomics, 19, 624.

22. Crowley, L.M. (2025) The genome sequence of a chironomid fly, Chironomus tentans Fabricius, 1805. Wellcome Open Res, 10, 133.

23. Pettrich, L.C., King, R., Field, L.M. and Waldvogel, A.-M. (2025) High-quality genome assembly of Chironomus riparius and its population history in European populations. G3 Genes|Genomes|Genetics, 15.

24. Holt, R.A., Subramanian, G.M., Halpern, A., Sutton, G.G., Charlab, R., Nusskern, D.R., Wincker, P., Clark, A.G., Ribeiro, J.M., Wides, R. et al. (2002) The genome sequence of the malaria mosquito Anopheles gambiae. Science, 298, 129–149.

25. Nene, V., Wortman, J.R., Lawson, D., Haas, B., Kodira, C., Tu, Z.J., Loftus, B., Xi, Z., Megy, K., Grabherr, M. et al. (2007) Genome sequence of Aedes aegypti, a major arbovirus vector. Science, 316, 1718–1723.

26. dos Santos, G., Schroeder, A.J., Goodman, J.L., Strelets, V.B., Crosby, M.A., Thurmond, J., Emmert, D.B., Gelbart, W.M. and Consortium, t.F. (2015) FlyBase: introduction of the Drosophila melanogaster Release 6 reference genome assembly and large-scale migration of genome annotations. Nucleic Acids Res., 43, D690–D697.

27. Oosterbroek, P. and Courtney, G. (1995) Phylogeny of the nematocerous families of Diptera (Insecta). Zoological Journal of the Linnean Society, 115, 267–311.

28. Wiegmann, B.M., Trautwein, M.D., Winkler, I.S., Barr, N.B., Kim, J.-W., Lambkin, C., Bertone, M.A., Cassel, B.K., Bayless, K.M., Heimberg, A.M. et al. (2011) Episodic radiations in the fly tree of life. Proceedings of the National Academy of Sciences, 108, 5690–5695.

29. Nell, L.A., Weng, Y.-M., Phillips, J.S., Botsch, J.C., Book, K.R., Einarsson, Á., Ives, A.R. and Schoville, S.D. (2024) Shared Features Underlying Compact Genomes and Extreme Habitat Use in Chironomid Midges. Genome Biol. Evol., 16.

30. Zhu, X., Pruess, K.P. and Powers, T.O. (1998) Mitochondrial DNA polymorphism in a black fly, Simulium vittatum (Diptera: Simuliidae). Can. J. Zool., 76, 440–447.

31. Adrien Leger, T.L. (2019) pycoQC, interactive quality control for Oxford Nanopore Sequencing. Journal of Open Source Software, 4.

32. De Coster, W., D’Hert, S., Schultz, D.T., Cruts, M. and Van Broeckhoven, C. (2018) NanoPack: visualizing and processing long-read sequencing data. Bioinformatics, 34, 2666–2669.

33. Wood, D.E. and Salzberg, S.L. (2014) Kraken: ultrafast metagenomic sequence classification using exact alignments. Genome Biol., 15, R46.

34. Cheng, H., Concepcion, G.T., Feng, X., Zhang, H. and Li, H. (2021) Haplotype-resolved de novo assembly using phased assembly graphs with hifiasm. Nature Methods, 18, 170–175.

35. Durand, N.C., Robinson, J.T., Shamim, M.S., Machol, I., Mesirov, J.P., Lander, E.S. and Aiden, E.L. (2016) Juicebox Provides a Visualization System for Hi-C Contact Maps with Unlimited Zoom. Cell Syst, 3, 99–101.

36. Gurevich, A., Saveliev, V., Vyahhi, N. and Tesler, G. (2013) QUAST: quality assessment tool for genome assemblies. Bioinformatics, 29, 1072–1075.

37. Cabanettes, F. and Klopp, C. (2018) D-GENIES: dot plot large genomes in an interactive, efficient and simple way. PeerJ, 6, e4958.

38. Flynn, J.M., Hubley, R., Goubert, C., Rosen, J., Clark, A.G., Feschotte, C. and Smit, A.F. (2020) RepeatModeler2 for automated genomic discovery of transposable element families. Proc Natl Acad Sci U S A, 117, 9451–9457.

39. Hoede, C., Arnoux, S., Moisset, M., Chaumier, T., Inizan, O., Jamilloux, V. and Quesneville, H. (2014) PASTEC: an automatic transposable element classification tool. PLoS One, 9, e91929.

40. Bao, W., Kojima, K.K. and Kohany, O. (2015) Repbase Update, a database of repetitive elements in eukaryotic genomes. Mob DNA, 6, 11.

41. Goubert, C., Craig, R.J., Bilat, A.F., Peona, V., Vogan, A.A. and Protasio, A.V. (2022) A beginner’s guide to manual curation of transposable elements. Mob DNA, 13, 7.

42. Storer, J., Hubley, R., Rosen, J., Wheeler, T.J. and Smit, A.F. (2021) The Dfam community resource of transposable element families, sequence models, and genome annotations. Mob DNA, 12, 2.

43. Marchler-Bauer, A., Derbyshire, M.K., Gonzales, N.R., Lu, S., Chitsaz, F., Geer, L.Y., Geer, R.C., He, J., Gwadz, M., Hurwitz, D.I. et al. (2015) CDD: NCBI’s conserved domain database. Nucleic Acids Res., 43, D222–226.

44. Tarailo-Graovac, M. and Chen, N. (2009) Using RepeatMasker to Identify Repetitive Elements in Genomic Sequences. Current Protocols, 25.**1**, 4–10.

45. Quinlan, A.R. and Hall, I.M. (2010) BEDTools: a flexible suite of utilities for comparing genomic features. Bioinformatics, 26, 841–842.

46. Martin, M. (2011) Cutadapt removes adapter sequences from high-throughput sequencing reads. EMBnet Journal.

47. Dobin, A., Davis, C.A., Schlesinger, F., Drenkow, J., Zaleski, C., Jha, S., Batut, P., Chaisson, M. and Gingeras, T.R. (2013) STAR: ultrafast universal RNA-seq aligner. Bioinformatics, 29, 15–21.

48. Grabherr, M.G., Haas, B.J., Yassour, M., Levin, J.Z., Thompson, D.A., Amit, I., Adiconis, X., Fan, L., Raychowdhury, R., Zeng, Q. et al. (2011) Full-length transcriptome assembly from RNA-Seq data without a reference genome. Nat. Biotechnol., 29, 644–652.

49. Consortium, T.U. (2015) UniProt: a hub for protein information. Nucleic Acids Res., 43, D204–D212.

50. Holt, C. and Yandell, M. (2011) MAKER2: an annotation pipeline and genome-database management tool for second-generation genome projects. BMC Bioinformatics, 12, 491.

51. Korf, I. (2004) Gene finding in novel genomes. BMC Bioinformatics, 5, 59.

52. Holst, F., Bolger, A.M., Kindel, F., Günther, C., Maß, J., Triesch, S., Kiel, N., Saadat, N., Ebenhöh, O., Usadel, B. et al. (2026) Helixer: ab initio prediction of primary eukaryotic gene models combining deep learning and a hidden Markov model. Nat Methods, 23, 732–739.

53. Batut, B., Hiltemann, S., Bagnacani, A., Baker, D., Bhardwaj, V., Blank, C., Bretaudeau, A., Brillet-Guéguen, L., Čech, M., Chilton, J. et al. (2018) Community-Driven Data Analysis Training for Biology. Cell Systems, 6, 752–758.e751.

54. Hiltemann, S., Rasche, H., Gladman, S., Hotz, H.R., Larivière, D., Blankenberg, D., Jagtap, P.D., Wollmann, T., Bretaudeau, A., Goué, N. et al. (2023) Galaxy Training: A powerful framework for teaching! *PLoS Comput*. Biol., 19, e1010752.

55. Pertea, G. and Pertea, M. (2020) GFF Utilities: GffRead and GffCompare. F1000Res, 9.

56. Skyler Kuhn, M.T. (2024). Zenodo, Vol. 2024.

57. Wingett, S.W. and Andrews, S. (2018) FastQ Screen: A tool for multi-genome mapping and quality control. F1000Res, 7, 1338.

58. Wood, D.E., Lu, J. and Langmead, B. (2019) Improved metagenomic analysis with Kraken 2. Genome Biol, 20, 257.

59. Garcia-Alcalde, F., Okonechnikov, K., Carbonell, J., Cruz, L.M., Gotz, S., Tarazona, S., Dopazo, J., Meyer, T.F. and Conesa, A. (2012) Qualimap: evaluating next-generation sequencing alignment data. Bioinformatics, 28, 2678–2679.

60. Wang, L., Wang, S. and Li, W. (2012) RSeQC: quality control of RNA-seq experiments. Bioinformatics, 28, 2184–2185.

61. Li, B. and Dewey, C.N. (2011) RSEM: accurate transcript quantification from RNA-Seq data with or without a reference genome. BMC Bioinformatics, 12, 323.

62. Love, M.I., Huber, W. and Anders, S. (2014) Moderated estimation of fold change and dispersion for RNA-seq data with DESeq2. Genome Biol., 15, 550.

63. Zheng, Z., Li, S., Su, J., Leung, A.W., Lam, T.W. and Luo, R. (2022) Symphonizing pileup and full-alignment for deep learning-based long-read variant calling. Nat Comput Sci, 2, 797–803.

64. Martin, M., Ebert, P. and Marschall, T. (2023) Read-Based Phasing and Analysis of Phased Variants with WhatsHap. Methods Mol. Biol., 2590, 127–138.

65. Rausch, T., Zichner, T., Schlattl, A., Stütz, A.M., Benes, V. and Korbel, J.O. (2012) DELLY: structural variant discovery by integrated paired-end and split-read analysis. Bioinformatics, 28, i333–i339.

66. Gladyshev, E.A., Meselson, M. and Arkhipova, I.R. (2008) Massive horizontal gene transfer in bdelloid rotifers. Science, 320, 1210–1213.

67. Boschetti, C., Carr, A., Crisp, A., Eyres, I., Wang-Koh, Y., Lubzens, E., Barraclough, T.G., Micklem, G. and Tunnacliffe, A. (2012) Biochemical diversification through foreign gene expression in bdelloid rotifers. PLoS Genet., 8, e1003035.

68. Donath, A., Jühling, F., Al-Arab, M., Bernhart, S.H., Reinhardt, F., Stadler, P.F., Middendorf, M. and Bernt, M. (2019) Improved annotation of protein-coding genes boundaries in metazoan mitochondrial genomes. Nucleic Acids Res., 47, 10543–10552.

69. Bernt, M., Donath, A., Jühling, F., Externbrink, F., Florentz, C., Fritzsch, G., Pütz, J., Middendorf, M. and Stadler, P.F. (2013) MITOS: improved de novo metazoan mitochondrial genome annotation. Mol. Phylogenet. Evol., 69, 313–319.

70. Katoh, K., Misawa, K., Kuma, K.i. and Miyata, T. (2002) MAFFT: a novel method for rapid multiple sequence alignment based on fast Fourier transform. Nucleic Acids Res., 30, 3059–3066.

71. Cock, P.J., Antao, T., Chang, J.T., Chapman, B.A., Cox, C.J., Dalke, A., Friedberg, I., Hamelryck, T., Kauff, F., Wilczynski, B. and de Hoon, M.J. (2009) Biopython: freely available Python tools for computational molecular biology and bioinformatics. Bioinformatics, 25, 1422–1423.

72. Ribeiro, J.M.C. and Arcà, B. (2009), Adv. In Insect Phys. Academic Press, Vol. 37, pp. 59–118.

73. Martin-Martin, I., Paige, A., Valenzuela Leon, P.C., Gittis, A.G., Kern, O., Bonilla, B., Chagas, A.C., Ganesan, S., Smith, L.B., Garboczi, D.N. and Calvo, E. (2020) ADP binding by the Culex quinquefasciatus mosquito D7 salivary protein enhances blood feeding on mammals. Nature Communications, 11, 2911.

74. Martin-Martin, I., Kern, O., Brooks, S., Smith, L.B., Valenzuela-Leon, P.C., Bonilla, B., Ackerman, H. and Calvo, E. (2021) Biochemical characterization of AeD7L2 and its physiological relevance in blood feeding in the dengue mosquito vector, Aedes aegypti. Febs j, 288, 2014–2029.

75. Feyereisen, R. (2012) In Gilbert, L. I. (ed.), Insect Molecular Biology and Biochemistry. Academic Press, San Diego, pp. 236–316.

76. Carey, A.F. and Carlson, J.R. (2011) Insect olfaction from model systems to disease control. Proc. Natl. Acad. Sci. U. S. A., 108, 12987–12995.

77. Willis, J.H. (2010) Structural cuticular proteins from arthropods: annotation, nomenclature, and sequence characteristics in the genomics era. Insect Biochem. Mol. Biol., 40, 189–204.

78. Kanost, M.R. and Jiang, H. (2015) Clip-domain serine proteases as immune factors in insect hemolymph. Curr Opin Insect Sci, 11, 47–55.

79. Wojda, I., Cytryńska, M., Zdybicka-Barabas, A. and Kordaczuk, J. (2020) Insect Defense Proteins and Peptides. Subcell. Biochem., 94, 81–121.

80. Manniello, M.D., Moretta, A., Salvia, R., Scieuzo, C., Lucchetti, D., Vogel, H., Sgambato, A. and Falabella, P. (2021) Insect antimicrobial peptides: potential weapons to counteract the antibiotic resistance. Cell. Mol. Life Sci., 78, 4259–4282.

81. Papanicolaou, A., Woo, A., Brei, B., Ma, D., Masedunskas, A., Gray, E., Xiao, G.G., Cho, S. and Brockhouse, C. (2013) Novel aquatic silk genes Simulium (Psilozia) vittatum (Zett) Diptera: Simuliidae. Insect Biochem. Mol. Biol., 43, 1181–1188.

82. Gries, J., Ebdon, S., Bliznina, A., Collins, J., Hodson, C.N., Mathers, T.C., Maulana, A., McCarthy, S.A., Paulini, M., Absolon, D.E. et al. (2026) 340 dipteran genomes reveal the origin of Muller elements and sex chromosomes in Diptera. bioRxiv, 2026.2006.2001.729285.

83. Iturbe-Ormaetxe, I., Burke, G.R., Riegler, M. and O’Neill, S.L. (2005) Distribution, expression, and motif variability of ankyrin domain genes in Wolbachia pipientis. J. Bacteriol., 187, 5136–5145.

84. Siozios, S., Ioannidis, P., Klasson, L., Andersson, S.G., Braig, H.R. and Bourtzis, K. (2013) The diversity and evolution of Wolbachia ankyrin repeat domain genes. PLoS One, 8, e55390.

85. Sproul, J.S., Hotaling, S., Heckenhauer, J., Powell, A., Marshall, D., Larracuente, A.M., Kelley, J.L., Pauls, S.U. and Frandsen, P.B. (2023) Analyses of 600+ insect genomes reveal repetitive element dynamics and highlight biodiversity-scale repeat annotation challenges. Genome Res., 33, 1708–1717.

86. Woodford, L., Bianco, G., Ivanova, Y., Dale, M., Elmer, K., Rae, F., Larcombe, S.D., Helm, B., Ferguson, H.M. and Baldini, F. (2018) Vector species-specific association between natural Wolbachia infections and avian malaria in black fly populations. Sci Rep, 8, 4188.

