## Supplemental Note for "*Chromosome-Level Genome of Simulium vittatum* Links Black Fly Cytogenetics to Genome Organization and Evolution"

### Supplementary Material

**Supplementary Figure 1**

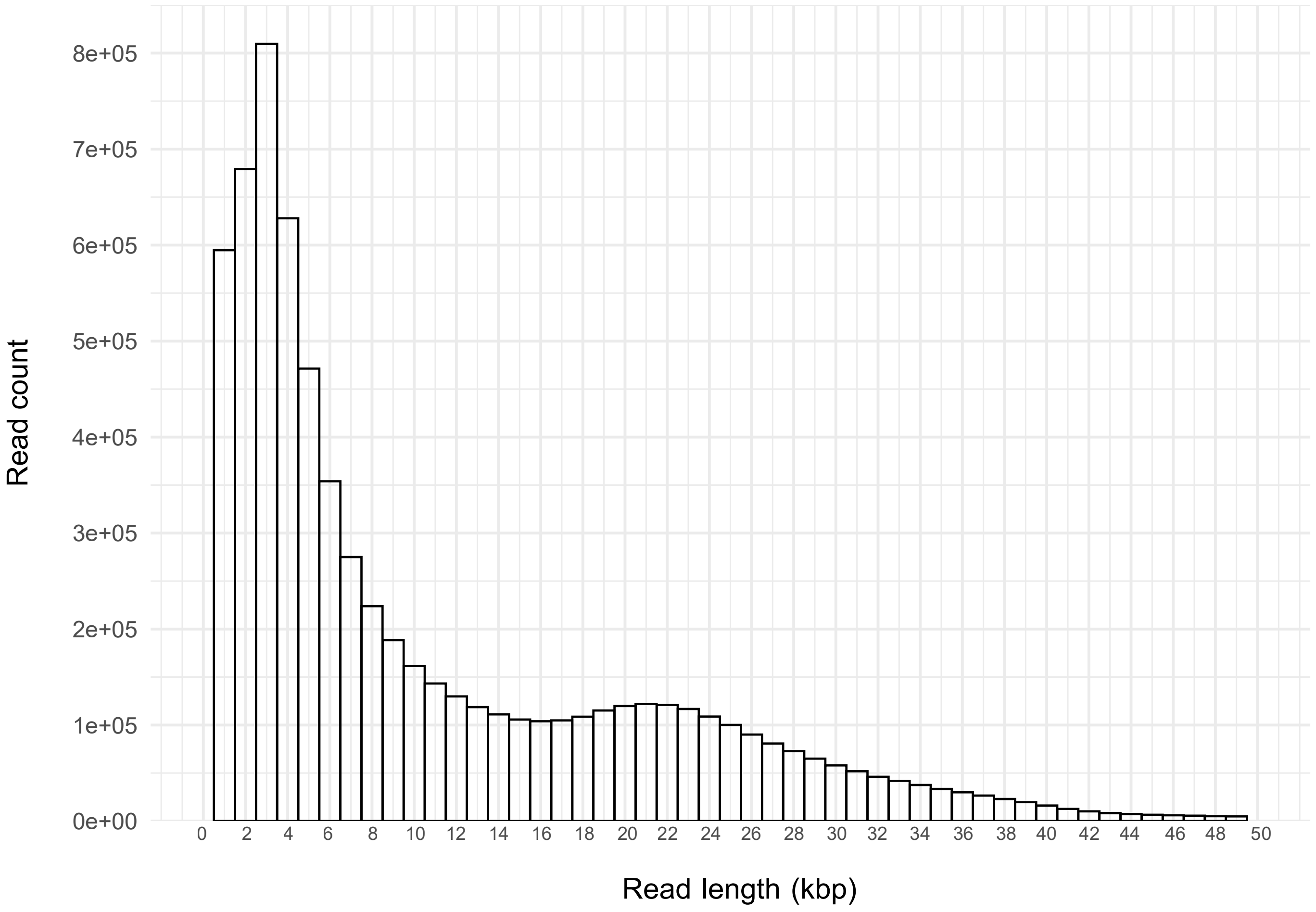


*Figure S1: Distribution of Oxford Nanopore long-read lengths used for genome assembly. The visualization is truncated at 50 kb for clarity.*

*Suppl. Table S2: Distribution of long-read sequencing lengths for the Simulium vittatum genome, summarized in 1 kb bins.*

| Read length interval (bp) | Read count | Percentage of reads (%) |
|---|---|---|
| 0-999 | 778008 | 10.57 |
| 1000-1999 | 543619 | 7.39 |
| 2000-2999 | 809356 | 11.00 |
| 3000-3999 | 726413 | 9.87 |
| 4000-4999 | 542887 | 7.38 |
| 5000-5999 | 408802 | 5.55 |
| 6000-6999 | 309481 | 4.20 |
| 7000-7999 | 247144 | 3.36 |
| 8000-8999 | 204668 | 2.78 |
| 9000-9999 | 174202 | 2.37 |
| 10000-10999 | 151672 | 2.06 |
| 11000-11999 | 136091 | 1.85 |
| 12000-12999 | 123808 | 1.68 |
| 13000-13999 | 114877 | 1.56 |
| 14000-14999 | 107658 | 1.46 |
| 15000-15999 | 104913 | 1.43 |
| 16000-16999 | 103718 | 1.41 |
| 17000-17999 | 106217 | 1.44 |
| 18000-18999 | 111986 | 1.52 |
| 19000-19999 | 117739 | 1.60 |
| 20000-20999 | 121175 | 1.65 |
| 21000-21999 | 122370 | 1.66 |
| 22000-22999 | 119293 | 1.62 |
| 23000-23999 | 112876 | 1.53 |
| 24000-24999 | 104850 | 1.42 |
| 25000-25999 | 94886 | 1.29 |
| 26000-26999 | 85308 | 1.16 |
| 27000-27999 | 76871 | 1.04 |
| 28000-28999 | 68870 | 0.94 |
| 29000-29999 | 61062 | 0.83 |
| 30000-30999 | 55039 | 0.75 |
| 31000-31999 | 48888 | 0.66 |
| 32000-32999 | 44033 | 0.60 |
| 33000-33999 | 39628 | 0.54 |
| 34000-34999 | 35733 | 0.49 |
| 35000-35999 | 31350 | 0.43 |
| 36000-36999 | 28110 | 0.38 |
| 37000-37999 | 24982 | 0.34 |
| 38000-38999 | 21155 | 0.29 |
| 39000-39999 | 17904 | 0.24 |
| 40000-40999 | 14321 | 0.19 |
| 41000-41999 | 11423 | 0.16 |
| 42000-42999 | 9016 | 0.12 |

Suppl. Table S2. Continued

| Read length interval (bp) | Read count | Percentage of reads (%) |
|---|---|---|
| 43000-43999 | 7632 | 0.10 |
| 44000-44999 | 6901 | 0.09 |
| 45000-45999 | 6273 | 0.09 |
| 46000-46999 | 5830 | 0.08 |
| 47000-47999 | 5247 | 0.07 |
| 48000-48999 | 4985 | 0.07 |
| 49000-49999 | 4579 | 0.06 |
| 50000-50999 | 4317 | 0.06 |
| 51000-51999 | 3830 | 0.05 |
| 52000-52999 | 3645 | 0.05 |
| 53000-53999 | 3379 | 0.05 |
| 54000-54999 | 3063 | 0.04 |
| 55000-55999 | 2810 | 0.04 |
| 56000-56999 | 2505 | 0.03 |
| 57000-57999 | 2390 | 0.03 |
| 58000-58999 | 2176 | 0.03 |
| 59000-59999 | 2015 | 0.03 |
| 60000-60999 | 1737 | 0.02 |
| 61000-61999 | 1543 | 0.02 |
| 62000-62999 | 1407 | 0.02 |
| 63000-63999 | 1236 | 0.02 |
| 64000-64999 | 1175 | 0.02 |
| 65000-65999 | 1031 | 0.01 |
| 66000-66999 | 904 | 0.01 |
| 67000-67999 | 764 | 0.01 |
| 68000-68999 | 667 | 0.01 |
| 69000-69999 | 609 | 0.01 |
| 70000-70999 | 535 | 0.01 |
| 71000-71999 | 495 | 0.01 |
| 72000-72999 | 400 | 0.01 |
| 73000-73999 | 397 | 0.01 |
| 74000-74999 | 305 | 0.00 |
| 75000-75999 | 262 | 0.00 |
| 76000-76999 | 239 | 0.00 |
| 77000-77999 | 246 | 0.00 |
| 78000-78999 | 178 | 0.00 |
| 79000-79999 | 224 | 0.00 |
| 80000-80999 | 165 | 0.00 |
| 81000-81999 | 152 | 0.00 |
| 82000-82999 | 122 | 0.00 |
| 83000-83999 | 127 | 0.00 |
| 84000-84999 | 84 | 0.00 |
| 85000-85999 | 96 | 0.00 |
| 86000-86999 | 85 | 0.00 |
| 87000-87999 | 65 | 0.00 |
| 88000-88999 | 59 | 0.00 |

Suppl. Table S2. Continued

| Read length interval (bp) | Read count | Percentage of reads (%) |
|---|---|---|
| 89000-89999 | 60 | 0.00 |
| 90000-90999 | 66 | 0.00 |
| 91000-91999 | 68 | 0.00 |
| 92000-92999 | 48 | 0.00 |
| 93000-93999 | 52 | 0.00 |
| 94000-94999 | 42 | 0.00 |
| 95000-95999 | 34 | 0.00 |
| 96000-96999 | 29 | 0.00 |
| 97000-97999 | 36 | 0.00 |
| 98000-98999 | 23 | 0.00 |
| 99000-99999 | 25 | 0.00 |
| 1e+05-100999 | 27 | 0.00 |
| 101000-101999 | 30 | 0.00 |
| 102000-102999 | 19 | 0.00 |
| 103000-103999 | 20 | 0.00 |
| 104000-104999 | 16 | 0.00 |
| 105000-105999 | 20 | 0.00 |
| 106000-106999 | 8 | 0.00 |
| 107000-107999 | 16 | 0.00 |
| 108000-108999 | 9 | 0.00 |
| 109000-109999 | 14 | 0.00 |
| 110000-110999 | 12 | 0.00 |
| 111000-111999 | 11 | 0.00 |
| 112000-112999 | 7 | 0.00 |
| 113000-113999 | 8 | 0.00 |
| 114000-114999 | 4 | 0.00 |
| 115000-115999 | 10 | 0.00 |
| 116000-116999 | 6 | 0.00 |
| 117000-117999 | 5 | 0.00 |
| 118000-118999 | 4 | 0.00 |
| 119000-119999 | 3 | 0.00 |
| 120000-120999 | 5 | 0.00 |
| 121000-121999 | 14 | 0.00 |
| 122000-122999 | 12 | 0.00 |
| 123000-123999 | 3 | 0.00 |
| 124000-124999 | 4 | 0.00 |
| 125000-125999 | 1 | 0.00 |
| 126000-126999 | 6 | 0.00 |
| 127000-127999 | 4 | 0.00 |
| 128000-128999 | 6 | 0.00 |
| 129000-129999 | 2 | 0.00 |
| 130000-130999 | 4 | 0.00 |
| 131000-131999 | 3 | 0.00 |
| 132000-132999 | 4 | 0.00 |
| 133000-133999 | 2 | 0.00 |
| 134000-134999 | 4 | 0.00 |

Suppl. Table S2. Continued

| Read length interval (bp) | Read count | Percentage of reads (%) |
|---|---|---|
| 135000-135999 | 3 | 0.00 |
| 136000-136999 | 1 | 0.00 |
| 137000-137999 | 1 | 0.00 |
| 140000-140999 | 3 | 0.00 |
| 141000-141999 | 1 | 0.00 |
| 143000-143999 | 1 | 0.00 |
| 144000-144999 | 2 | 0.00 |
| 145000-145999 | 1 | 0.00 |
| 146000-146999 | 1 | 0.00 |
| 148000-148999 | 1 | 0.00 |
| 151000-151999 | 3 | 0.00 |
| 156000-156999 | 1 | 0.00 |
| 161000-161999 | 1 | 0.00 |
| 163000-163999 | 1 | 0.00 |
| 180000-180999 | 1 | 0.00 |
| 183000-183999 | 1 | 0.00 |
| 184000-184999 | 1 | 0.00 |
| 197000-197999 | 1 | 0.00 |
| 223000-223999 | 1 | 0.00 |
| 271000-271999 | 1 | 0.00 |
| 329000-329999 | 1 | 0.00 |

#### Supplementary Note S1. Genomic and transcriptomic characterization of larval silk gland proteins

##### Identification of previously characterized larval silk gland protein genes

Black fly larvae use silk for attachment and movement on submerged substrates and for construction of the pupal cocoon. Three major *Simulium vittatum* silk gland proteins, SGP-1, SGP-2, and SGP-3, were previously characterized (1). The published protein sequences AHF71320.1, AHF71321.1, and AHF71322.1 were used as queries to identify the corresponding loci in the new *S. vittatum* genome and transcriptome. SGP1 corresponded to SV00009995-RA, SGP-2 to SV00009996-RA, and SGP-3 to SV00009306-RA (Supplementary Alignments S1–S3). All three proteins contained N-terminal signal peptides predicted by SignalP, and functional annotation identified the corresponding published silk protein as the best Diptera match (Supplementary Table S4).

Developmental transcript profiles were consistent with the established roles of these proteins in larval silk production. SGP-1, SGP-2, and SGP-3 showed larval transcript abundances of 18,317.75, 20,560.98, and 527.26 TPM, respectively, with markedly lower abundance in pupal and adult samples (Supplementary Table S3). Because each developmental condition was represented by one pooled biological sample, these values are presented descriptively and were not subjected to statistical differential-expression testing.

##### Genomic organization of the SGP loci

All three loci occur on chromosome II. SGP-1 (SV00009995-RA; 100,460,812–100,463,044 bp) and SGP-2 (SV00009996-RA; 100,466,099–100,467,495 bp) are closely linked, with 3,054 bp separating their annotated transcript boundaries, and both are oriented on the minus strand. SGP-3 (SV00009306-RA; 86,972,727–86,973,853 bp) occurs approximately 13.5 Mb from the SGP-1/SGP-2 region, also on the minus strand (Suppl. Table S9). The close physical association of SGP-1 and SGP-2 is consistent with local duplication, although their evolutionary relationship was not investigated here.

##### Transcript-supported reconstruction of SGP2

SGP-2 required additional examination. The genome-derived SV00009996-RA model predicts a 365-aa protein that is highly conserved relative to the previously reported 336-aa SGP-2 through residue 128 (123/128 identical residues; 96.1% identity) but diverges abruptly thereafter. Examination of the gene structure showed that the reading-frame discrepancy coincides with the boundary between the two annotated coding exons. The first CDS segment spans 100,466,969-100,467,352 bp and contains exactly 384 nt, corresponding to the first 128 codons, whereas the second CDS segment spans 100,466,187-100,466,900 bp. The gene is encoded on the minus strand.

Incorporation of a single G immediately after CDS nucleotide 384 in the reconstructed sequence restores the downstream reading frame and produces a 336-aa protein with 97.0% identity and 98.2% similarity to the previously reported SGP2 (Supplementary Alignment S2).

Larval RNA-seq provided independent support for this reconstructed sequence. The genome-derived junction sequence TCATCTCACTCTTCTGATGGGAATCAGGAAAGGGCG was not detected as an exact match in either R1 or R2 reads. In contrast, the corresponding sequence containing the additional G, TCATCTCACTCTTCTGGATGGGAATCAGGAAAGGGCG, was detected in 13,369 R1 and 12,526 R2 reads. Individual 150-bp reads containing this sequence mapped specifically to the SGP2 genomic region: one representative read contained a 105-bp segment with 100% identity to chromosome II positions 100,467,071-100,466,967 and a 47-bp segment with 100% identity to positions 100,466,901-100,466,855, consistent with an SGP2 transcript spanning the annotated intron.

The genomic region surrounding SGP2 was also compared between the final chromosome-scale assembly and the original ONT-derived contig assembly. A 1,001-bp region encompassing the SGP2 junction matched contig_1655 with 100% nucleotide identity, indicating that the genomic sequence discrepancy was already present in the underlying contig assembly and was not introduced during Hi-C scaffolding (Suppl. Table S10).

Together, the protein homology and larval RNA-seq evidence support a 336-aa SGP2 coding sequence containing the additional nucleotide at the exon junction. The available data do not establish whether the discrepancy between the assembled genomic sequence and expressed transcript reflects a consensus error in the underlying genome assembly, an annotation-related issue, or another source of genomic-transcript sequence difference. Independent genomic read-level or molecular validation will be required to resolve its origin.

#### Reference


1. Papanicolaou, A., Woo, A., Brei, B., Ma, D., Masedunskas, A., Gray, E., Xiao, G.G., Cho, S. and Brockhouse, C. (2013) Novel aquatic silk genes Simulium (Psilozia) vittatum (Zett) Diptera: Simuliidae. *Insect Biochem. Mol. Biol.*, **43**, 1181–1188.

#### Supplementary Methods: SGP2 sequence validation

The SGP2 CDS was reconstructed directly from the chromosome-scale reference using the annotated CDS coordinates for SV00009996-RA. Because the locus is encoded on the minus strand, genomic CDS segments were reverse-complemented and concatenated in transcript order. The resulting genome-derived CDS comprised 1,098 nt. Junction sequences representing the genome-derived model and a reconstructed sequence containing one additional G at the boundary between the two annotated coding exons were searched as exact nucleotide matches against paired-end larval RNA-seq reads. R1 and R2 were evaluated separately. Representative RNA reads containing the reconstructed junction were aligned against the chromosome-scale assembly using BLASTN to confirm their genomic origin. A 1,001-bp genomic region encompassing the SGP2 junction was additionally aligned against the original ONT-derived contig assembly to determine whether the discrepancy was introduced during chromosome scaffolding. Sequence manipulation and searches were performed using SeqKit v2.13.0, BLAST+ v2.15.0+, and SAMtools 1.23.

**Supplementary Table S9. Genomic and transcriptomic characteristics of previously characterized *S. vittatum* silk gland proteins.**

| Protein | NCBI Accession | Transcript | Chromosome | Coordinates | Strand | Published/New Protein | Identity | Similarity | Larval TPM |
|---|---|---|---|---|---|---|---|---|---|
| SGP1 | AHF71320.1 | SV00009995-RA | II | 100,460,812–100,463,044 | - | 527 / 530 aa | 97.9% | 98.5% | 18,317.75 |
| SGP2, genome derived | AHF71321.1 | SV00009996-RA | II | 100,466,099-100,467,495 | - | 336 / 365 aa | 96.1% | 97.7% | 20,560.98 |
| SGP2, RNA-supported reconstruction | AHF71321.1 | SV00009996-RA | II | Same locus | - | 336 / 336 aa | 97.0% | 98.2% | - |
| SGP3 | AHF71322.1 | SV00009306-RA | II | 86,972,727-86,973,853 | - | 146 / 146 aa | 100.0% | 100.0% | 527.26 |

*Identity and similarity for the genome-derived SGP-2 are calculated over the 128-aa high-confidence N-terminal alignment preceding the reading-frame divergence. The RNA-supported reconstruction differs from the genome-derived model by one additional G at the boundary between the two annotated coding exons. Transcript abundances are descriptive because developmental samples were not biologically replicated.

Supplementary Table S10. Evidence supporting the reconstructed SGP2 exon junction

| Evidence | Genome-derived junction | +G reconstructed junction |
|---|---|---|
| Junction Sequence | ...TCTTCTGATGGGA... | ...TCTTCTGGATGGGA... |
| Difference | - | Additional G |
| Predicted protein length | 365 aa | 336 aa |
| Agreement with published SGP-2 | 96.1% identity over aa 1-128; diverges thereafter | 97.0% identity over 336 aa |
| Protein similarity | 97.7% over aa 1-128 | 98.2% similarity over 336 aa |
| Exact larval R1 matches | 0 | 13,369 |
| Exact larval R2 matches | 0 | 12,526 |
| Representative RNA read | - | Two 100%-identity segments mapping across SGP2 intron |
| Original ONT contig vs Final assembly | Same genome-derived sequence | - |

R1 and R2 counts are reported separately because they represent paired-end mates from the same sequencing library and should not be interpreted as independent biological replicates.

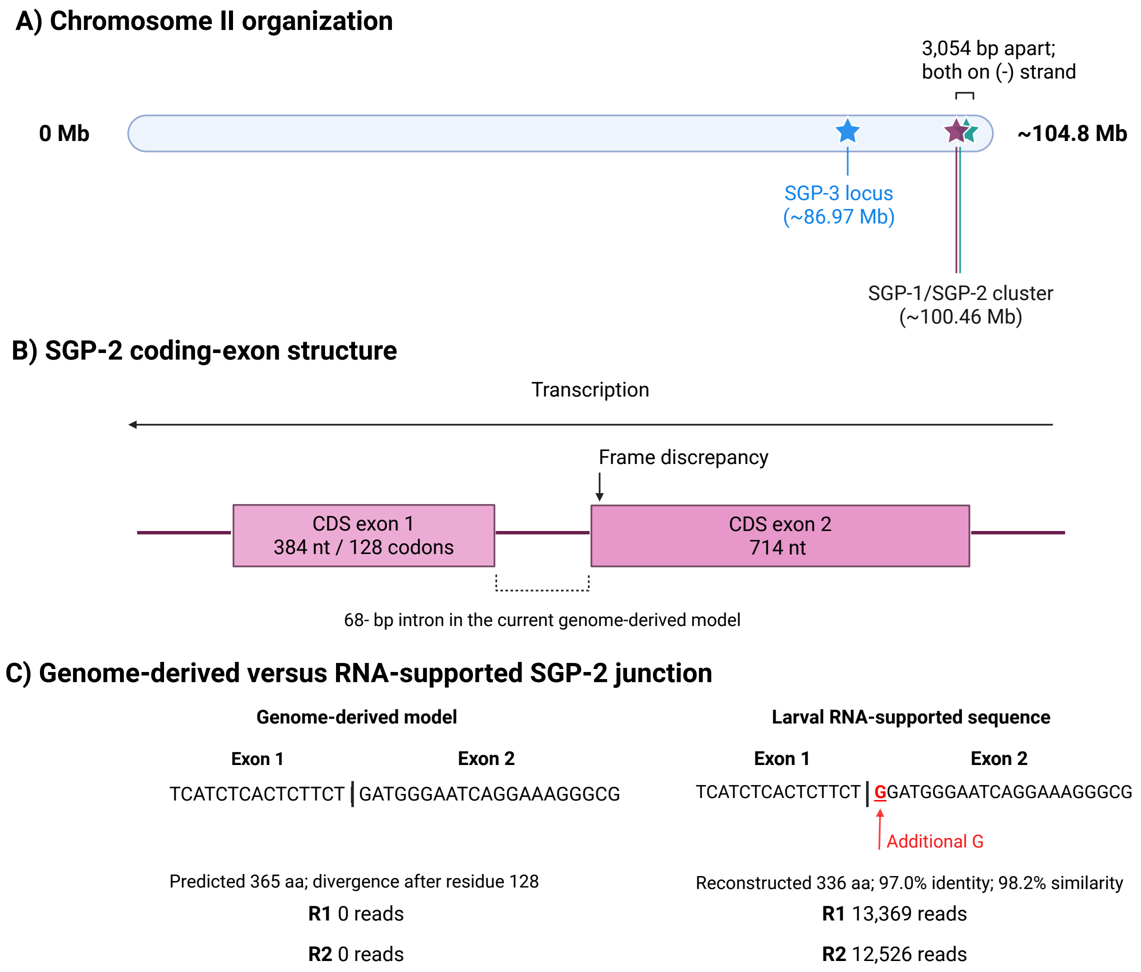


**Supplementary Figure S2: Genomic organization of *S. vittatum* silk gland protein genes and transcript-supported reconstruction of SGP2. (A)** Chromosomal locations of SGP1–SGP3 on chromosome II. SGP1 and SGP2 are separated by 3,054 bp between their annotated transcript boundaries and are both encoded on the minus strand; SGP3 occurs approximately 13.5 Mb from the SGP1/SGP2 region. **(B)** Annotated SGP2 gene structure. The first coding segment comprises 384 nt (128 codons), and the reading-frame discrepancy relative to previously characterized SGP2 occurs at the boundary between the two annotated coding exons. **(C)** Comparison of the genome-derived and RNA-supported junction sequences. Larval RNA reads support an additional G relative to the genome-derived model. The RNA-supported sequence was detected in 13,369 R1 and 12,526 R2 reads, whereas no exact matches to the corresponding genome-derived junction sequence were detected. Incorporation of the additional nucleotide restores a 336-aa protein with 97.0% identity and 98.2% similarity to previously characterized SGP2 (AHF71321.1). Created in BioRender. Sayuri Nishiduka Costa, E. (2026) https://BioRender.com/xtfkrm3

#### Supplementary Alignment S1 - SGP1

**Published SGP1 (AHF71320.1) versus the protein encoded by SV00009995-RA.** The sequences show 97.9% amino acid identity and 98.5% similarity. Vertical bars indicate identical residues and colons indicate conservative substitutions with positive BLOSUM62 scores.

```
AHF71320.1        1 MQSKILILLLVVVAINLVSSGVAPKKYRKGHYVGGYGKGYGKGYGYGKGYGKGYGY----   56
                    ||||||||||||||||||||||||||| ||||||||||||||||||||||||||||
SV00009995-RA     1 MQSKILILLLVVVAINLVSSGVAPKKYSKGHYVGGYGKGYGKGYGYGKGYGKGYGYGKGY   60
AHF71320.1       57 --GKGYGKGYGSGSGSAEYRRAIYAGNASGESGRAFAGYRGAGYDGGSSEEFGGRYMQAR  114
                      ||||||| |||||||||||||||||||||||||||||||||||||||||| |||||||
SV00009995-RA    61 GHGKGYGKGRGSGSGSAEYRRAIYAGNASGESGRAFAGYRGAGYDGGSSEEFSGRYMQAR  120
AHF71320.1      115 SGKDGEFKAGGDRNRGGPRGSKVSQNRDLKLNQYIGKEGAGAHYSGAFKDRASAEGAAHT  174
                    ||||||||||||||||||||||||||||||||||||||||||||||||||||||||||||
SV00009995-RA   121 SGKDGEFKAGGDRNRGGPRGSKVSQNRDLKLNQYIGKEGAGAHYSGAFKDRASAEGAAHT  180
AHF71320.1      175 SGERGDRQKWAASARASAEQHARQEAGSKERSMSASGARASSEAYKRGSGEKYARQQEQE  234
                    |||:|||||:|||||||||||||||| |||||||||||||||||||||||||||||||||
SV00009995-RA   181 SGEKGDRQKFAASARASAEQHARQEAASKERSMSASGARASSEAYKRGSGEKYARQQEQE  240
AHF71320.1      235 RARNAARAASRDSAEQHKRSRAESLEQHSGEKARQRSGWSRAQQSGESHERYEAARNARK  294
                    ||||||||||||||||||||||||||||||||||||||||||||||||||||||||||||
SV00009995-RA   241 RARNAARAASRDSAEQHKRSRAESLEQHSGEKARQRSGWSRAQQSGESHERYEAARNARK  300
AHF71320.1      295 RSNMRARNSREERARHSAEKMSRFRDMNAAEKGGAKAQKYGGENHQVYKYNGAMPGRKPI  354
                    ||||||||||||||||||||||||||||||||||||||||||||||||||||||||||||
SV00009995-RA   301 RSNMRARNSREERARHSAEKMSRFRDMNAAEKGGAKAQKYGGENHQVYKYNGAMPGRKPI  360
AHF71320.1      355 SIEAARNSLEKAQAKLELSSMESRGPRSFEKSMERMGRYSGQRLRKNYFGGSIEQAEGVA  414
                    ||||||||||||||||||||||||||||||||||||||||||||||||||||||||||| 
SV00009995-RA   361 SIEAARNSLEKAQAKLELSSMESRGPRSFEKSMERMGRYSGQRLRKNYFGGSIEQAEGVD  420
AHF71320.1      415 AYGKKTGPIDMQMVRNVQIKQKGDGIYDTLKRRGNYGYADGKGYYGKGYGYGKGYGKGHG  474
                    ||||||||||||||||| |||||||||||||||||||||||:||||||||||||||||||
SV00009995-RA   421 AYGKKTGPIDMQMVRNVNIKQKGDGIYDTLKRRGNYGYADGEGYYGKGYGYGKGYGKGHG  480
AHF71320.1      475 YGKGYGLGRYGRGYGYGKGWGKDYTVAPAAPVATTAAPATKPAVTAAPVTPCD  527
                    |||||| |||||||||||||||||||   ||||||||||| ||||||||||||
SV00009995-RA   481 YGKGYGHGRYGRGYGYGKGWGKDYTV---APVATTAAPATLPAVTAAPVTPCD  530
```

#### Supplementary Alignment S2 - SGP2 RNA-supported reconstruction

Published SGP2 (AHF71321.1) versus the RNA-supported reconstructed protein encoded at the SV00009996-RA locus. The reconstructed 336-aa sequence is derived from the junction sequenced containing the additional G supported by larval RNA-seq and shows 97.0% identity and 98.2% similarity to AHF71321.1.

```
AHF71321.1        1 MQSKTLLCFLVVLAISYATAGVPHRRYSDSCSDEHGRYVGGWGWGHGHGHGGHGWGKGRG   60
                    ||||||||||||||||||||||||||||||||||||  ||||||||||| ||:|||||||
SGP2-recon.       1 MQSKTLLCFLVVLAISYATAGVPHRRYSDSCSDEHGHNVGGWGWGHGHGAGGYGWGKGRG   60

AHF71321.1       61 YGGNVARYGGRRNFDLGRNEADKARSASWERVPMKRYQKYATLKPTKTAYNHVSPDGNVK  120
                    ||||||||||||||||||||||||||||||||||||||||||||||||||||||||||||
SGP2-recon.      61 YGGNVARYGGRRNFDLGRNEADKARSASWERVPMKRYQKYATLKPTKTAYNHVSPDGNVK  120

AHF71321.1      121 SWGSAHSSGWESGKGGAMEYHSGEKHDRYGGDRSKRYSGERRAYASGEDRSKQRAAAYRR  180
                    ||||:|||||||||||||||||||||||:|||||||||||||||||||||||||||||||
SGP2-recon.     121 SWGSSHSSGWESGKGGAMEYHSGEKHDRFGGDRSKRYSGERRAYASGEDRSKQRAAAYRR  180

AHF71321.1      181 SSGERKAAKQSTSKERSQGSRDTRRSAEAWAKNAGRSDESYSAERQRQASTEGHARQRKG  240
                    ||||||||||||||||||||||||||||||||||||||||||||||||||||||||||||
SGP2-recon.     181 SSGERKAAKQSTSKERSQGSRDTRRSAEAWAKNAGRSDESYSAERQRQASTEGHARQRKG  240

AHF71321.1      241 SGAQKRGAAYAKSAAEQAGYNKERETAKSSAERAARAASEGHRYDSGEKHARRSGEQNVK  300
                    |||||||||:|||||||||||||| |||||||||||||||||||||||||||||||||||
SGP2-recon.     241 SGAQKRGAAFAKSAAEQAGYNKERATAKSSAERAARAASEGHRYDSGEKHARRSGEQNVK  300

AHF71321.1      301 GSTERGARYATAHRGKFRDADGDSWSGEGYRPRNKH  336
                    ||||||||||||||||| ||| ||||||||||||||
SGP2-recon.     301 GSTERGARYATAHRGKFNDADADSWSGEGYRPRNKH  336
```

#### Supplementary Alignment S3 - SGP3

Published SGP3 (AHF71322.1) versus the protein encoded by SV00009306-RA. The proteins are identical across all 146 amino acids.

```
AHF71322.1        1 MQPKIIVCFLVVLTINLALASFAAPKGCKSIEITKENAAKPEKNSFEACARGFFARKKGG   60
                    ||||||||||||||||||||||||||||||||||||||||||||||||||||||||||||
SV00009306-RA     1 MQPKIIVCFLVVLTINLALASFAAPKGCKSIEITKENAAKPEKNSFEACARGFFARKKGG   60
AHF71322.1       61 AVGGWAGDWGRRRRGSGSASQESRGWGGRGWGKSRGYNRKGRGGYGKGRGGYGKRRGYYG  120
                    ||||||||||||||||||||||||||||||||||||||||||||||||||||||||||||
SV00009306-RA    61 AVGGWAGDWGRRRRGSGSASQESRGWGGRGWGKSRGYNRKGRGGYGKGRGGYGKRRGYYG  120
AHF71322.1      121 KGRGRYGKGRGYGNKYGKGKKYGRFY  146
                    ||||||||||||||||||||||||||
SV00009306-RA   121 KGRGRYGKGRGYGNKYGKGKKYGRFY  146
```
