## Supplemental Table 5 for "*Chromosome-Level Genome of Simulium vittatum* Links Black Fly Cytogenetics to Genome Organization and Evolution"

Table S5. Selected biologically relevant gene families identified in the S. vittatum genome and supporting secretion predictions.

| **Gene family** | **Predicted loci** | **SignalP SIG loci** | **Annotation-supported secreted loci** | **Dual-supported loci** |
| --- | --- | --- | --- | --- |
| CYP450 | 140 | 50 | 12 | 12 |
| Trypsin/Chymotrypsin | 137 | 105 | 118 | 99 |
| Cuticle | 133 | 115 | 26 | 26 |
| OR | 83 | 9 | 8 | 7 |
| GR | 65 | 13 | 10 | 9 |
| ABC | 53 | 4 | 4 | 4 |
| Chitin | 51 | 36 | 17 | 17 |
| OBP | 49 | 40 | 42 | 40 |
| Phospholipase | 42 | 24 | 19 | 19 |
| Peritrophin | 38 | 32 | 23 | 23 |
| GST | 29 | 2 | 0 | 0 |
| Cathepsin | 27 | 20 | 21 | 20 |
| IR | 25 | 14 | 7 | 7 |
| Carboxylesterase | 22 | 17 | 20 | 17 |
| CLIP | 21 | 17 | 18 | 15 |
| Juvenile hormone | 19 | 12 | 13 | 12 |
| HSP | 18 | 0 | 0 | 0 |
| Antigen 5 | 17 | 13 | 15 | 12 |
| Lysozyme | 10 | 8 | 1 | 1 |
| Ferritin | 8 | 3 | 1 | 1 |
| Vitellogenin | 8 | 6 | 0 | 0 |
| Aquaporin | 6 | 1 | 0 | 0 |
| Defensin | 5 | 1 | 1 | 1 |
| Apyrase | 4 | 4 | 4 | 4 |
| Cecropin | 1 | 0 | 0 | 0 |

**Notes:** Counts represent unique predicted loci after collapsing transcript isoforms and were based on the strongest available functional assignments from reference protein annotation and conserved-domain evidence. SignalP SIG loci are unique predicted loci with at least one transcript classified as SIG. Annotation-supported secreted loci are unique loci with at least one transcript assigned to a secretion-associated annotation category. Dual-supported loci satisfy both criteria. Gene families are not mutually exclusive; therefore, counts should not be summed across rows.

**Legends:** ABC, ATP-binding cassette transporter; CLIP, CLIP-domain serine protease; CYP450, cytochrome P450; GR, gustatory receptor; GST, glutathione S-transferase; HSP, heat-shock protein; IR, ionotropic receptor; OBP, odorant-binding protein; OR, odorant receptor.
