## Supplemental Table 7 for "*Chromosome-Level Genome of Simulium vittatum* Links Black Fly Cytogenetics to Genome Organization and Evolution"

Supplementary Table S7: Candidate symbiont-associated horizontally transferred genes identified in S. vittatum based on Alien Index (AI > 30) and HGT index (hU > 30). Best BLASTP hits are reported with accession numbers, broad taxonomic origin, source species, and predicted functional annotation.

| Protein ID | Best Hit ID | Broad Taxa | Species | Protein Annotation |
| --- | --- | --- | --- | --- |
| SV00006927-RA | WP_265016916.1 | Bacteria | Wolbachia sp. | Tetratricopeptide repeat protein |
| SV00006089-RA | KAJ6441863.1 | Fungi | Purpureocillium lavendulum | Hsp70 family chaperone |
| SV00004516-RA | WP_264705646.1 | Bacteria | Wolbachia sp. | Ankyrin repeat domain-containing protein |
| SV00006932-RA | WP_264377089.1 | Bacteria | Wolbachia sp. | Ankyrin repeat domain-containing protein |
| SV00010919-RA | KAL9584638.1 | Fungi | Teloschistes exilis | Hypothetical protein |
| SV00007412-RA | MDR0329725.1 | Bacteria | Rickettsia sp. | Hypothetical protein |
| SV00006931-RA | WP_019236511.1 | Bacteria | Wolbachia sp. | Tetratricopeptide repeat protein (partial) |
| SV00006930-RA | WP_410542226.1 | Bacteria | Wolbachia sp. | Tetratricopeptide repeat protein |
| SV00006929-RA | WP_264377910.1 | Bacteria | Wolbachia sp. | Ankyrin repeat domain-containing protein |
