## Supplemental Table 8 for "*Chromosome-Level Genome of Simulium vittatum* Links Black Fly Cytogenetics to Genome Organization and Evolution"

Suppl. Table S8: Overall Gene Composition and Features of the Simulium vittatum Mitochondrial Genome.

| **Molecule** | **Gene** | **Start** | **End** | **Orientation** |
| --- | --- | --- | --- | --- |
| PGA_scaffold_88__1_contigs__length_20965 | trnM(cat) | 1 | 70 | Forward |
| PGA_scaffold_88__1_contigs__length_20965 | nad2 | 100 | 964 | Forward |
| PGA_scaffold_88__1_contigs__length_20965 | trnW(tca) | 1111 | 1182 | Forward |
| PGA_scaffold_88__1_contigs__length_20965 | trnC(gca) | 1174 | 1238 | Reverse |
| PGA_scaffold_88__1_contigs__length_20965 | trnY(gta) | 1262 | 1327 | Reverse |
| PGA_scaffold_88__1_contigs__length_20965 | cox1 | 1340 | 2879 | Forward |
| PGA_scaffold_88__1_contigs__length_20965 | trnL2(taa) | 2874 | 2940 | Forward |
| PGA_scaffold_88__1_contigs__length_20965 | cox2 | 2992 | 3634 | Forward |
| PGA_scaffold_88__1_contigs__length_20965 | trnK(ctt) | 3634 | 3705 | Forward |
| PGA_scaffold_88__1_contigs__length_20965 | trnD(gtc) | 3723 | 3793 | Forward |
| PGA_scaffold_88__1_contigs__length_20965 | atp8 | 3793 | 3958 | Forward |
| PGA_scaffold_88__1_contigs__length_20965 | atp6 | 3957 | 4614 | Forward |
| PGA_scaffold_88__1_contigs__length_20965 | cox3 | 4644 | 5418 | Forward |
| PGA_scaffold_88__1_contigs__length_20965 | trnG(tcc) | 5426 | 5491 | Forward |
| PGA_scaffold_88__1_contigs__length_20965 | nad3 | 5542 | 5842 | Forward |
| PGA_scaffold_88__1_contigs__length_20965 | trnA(tgc) | 5856 | 5921 | Forward |
| PGA_scaffold_88__1_contigs__length_20965 | trnR(tcg) | 5923 | 5987 | Forward |
| PGA_scaffold_88__1_contigs__length_20965 | trnN(gtt) | 5990 | 6055 | Forward |
| PGA_scaffold_88__1_contigs__length_20965 | trnS1(gct) | 6055 | 6122 | Forward |
| PGA_scaffold_88__1_contigs__length_20965 | trnE(ttc) | 6122 | 6188 | Forward |
| PGA_scaffold_88__1_contigs__length_20965 | trnF(gaa) | 6202 | 6268 | Reverse |
| PGA_scaffold_88__1_contigs__length_20965 | nad5 | 6313 | 7894 | Reverse |
| PGA_scaffold_88__1_contigs__length_20965 | trnH(gtg) | 8005 | 8071 | Reverse |
| PGA_scaffold_88__1_contigs__length_20965 | nad4 | 8091 | 9369 | Reverse |
| PGA_scaffold_88__1_contigs__length_20965 | nad4l | 9407 | 9653 | Reverse |
| PGA_scaffold_88__1_contigs__length_20965 | trnT(tgt) | 9703 | 9770 | Forward |
| PGA_scaffold_88__1_contigs__length_20965 | trnP(tgg) | 9770 | 9835 | Reverse |
| PGA_scaffold_88__1_contigs__length_20965 | nad6 | 9873 | 10353 | Forward |
| PGA_scaffold_88__1_contigs__length_20965 | cob | 10370 | 11453 | Forward |
| PGA_scaffold_88__1_contigs__length_20965 | trnS2(tga) | 11498 | 11565 | Forward |
| PGA_scaffold_88__1_contigs__length_20965 | nad1 | 11612 | 12500 | Reverse |
| PGA_scaffold_88__1_contigs__length_20965 | trnL1(tag) | 12540 | 12606 | Reverse |
| PGA_scaffold_88__1_contigs__length_20965 | rrnL | 12610 | 13903 | Reverse |
| PGA_scaffold_88__1_contigs__length_20965 | trnV(tac) | 13945 | 14017 | Reverse |
| PGA_scaffold_88__1_contigs__length_20965 | rrnS | 14016 | 14807 | Reverse |
| PGA_scaffold_88__1_contigs__length_20965 | trnI(gat) | 15684 | 15752 | Forward |
| PGA_scaffold_88__1_contigs__length_20965 | trnQ(ttg) | 15757 | 15826 | Reverse |
